# The geometry and genetics of hybridization

**DOI:** 10.1101/862235

**Authors:** Hilde Schneemann, Bianca De Sanctis, Denis Roze, Nicolas Bierne, John J. Welch

## Abstract

We develop an analytical framework for predicting the fitness of hybrid genotypes, based on Fisher’s geometric model. We first show that all of the model parameters have a simple geometrical and biological interpretation. Hybrid fitness decomposes into intrinsic effects of hybridity and heterozygosity, and extrinsic measures of the (local) adaptedness of the parental lines; and all of these correspond to distances in a phenotypic space. We also show how these quantities change over the course of divergence, with convergence to a characteristic pattern of intrinsic isolation. Using individual-based simulations, we then show that the predictions apply to a wide range of population genetic regimes, and divergence conditions, including allopatry and parapatry, local adaptation and drift. We next connect our results to the quantitative genetics of line crosses in variable or patchy environments. This relates the geometrical distances to quantities that can be estimated from cross data, and provides a simple interpretation of the “composite effects” in the quantitative genetics partition. Finally, we develop extensions to the model, involving selectively-induced disequilibria, and variable phenotypic dominance. The geometry of fitness landscapes provides a unifying framework for understanding speciation, and wider patterns of hybrid fitness.

## Introduction

When genetically distinct populations meet and mate, their divergent alleles are brought together in new combinations. The fitness of these novel genotypes will influence the outcome of the hybridization, and might be a source of information about the divergence history of the populations (e.g., Demuth and Wade, 2005; Dobzhansky, 1937; Fraïsse et al., 2016; Gavrilets, 2004; Lynch, 1991; Rosas et al., 2010; Rundle and Whitlock, 2001; Welch, 2004; Yamaguchi and Otto, 2019).

To predict and interpret data from hybrids, various approaches have been used. One approach uses fitness landscapes (Dobzhansky, 1937; Gavrilets, 2004; Hill, 1982; Orr, 1995). Here, strong insights can come from simple models, with a few, biologically meaningful parameters; but such models often apply to a limited range of cases, and can be difficult to fit to data. A second approach uses quantitative genetics, applying to line crosses, statistical tools that were developed for single populations (Cockerham, 1980; Demuth and Wade, 2005; Hill, 1982; Lynch, 1991; Lynch and Walsh, 1998; Rundle and Whitlock, 2001, Chs. 9-10). This approach is fully general and widely applied, but the key quantities - the “composite effects” - can be difficult to interpret (with single populations, by contrast, a large body of theory can help us to understand the variance components; Barton, 2017; Hill et al., 2008; Mäki-Tanila and Hill, 2014; Walsh and Lynch, 2018).

Previous authors have gained fresh insights by combining these approaches (Demuth and Wade, 2005; Lynch, 1991; Yamaguchi and Otto, 2019). Here, following these authors, we draw an explicit connection between the quantitative genetics of line crosses (Hill, 1982; Rundle and Whitlock, 2001), and a class of fitness landscapes based on Fisher’s geometric model (Fisher, 1930, Ch. 2).

Fisher’s geometric model has been widely used to study single populations (Orr, 1998b; Hartl and Taubes, 1998; Walsh and Lynch, 2018, Ch. 27), and hybridization, and it can account for a large number of empirical patterns (Barton, 2001; Chevin et al., 2014; Fraïsse et al., 2016; Mani and Clarke, 1990; Rosas et al., 2010; Simon et al., 2018; Thompson et al., 2019; Yamaguchi and Otto, 2019). But despite these successes, some serious doubts remain. First, the model assumes that fitness is determined by a few quantitative traits, each with additive genetics. This assumption is less restrictive than it seems, because the “traits” need not be identified with standard quantitative traits (such as height or weight), but might emerge as an approximation to a variety of more complex phenotypic models, including many-to-one mappings (Fraïsse and Welch, 2019; Martin, 2014; Schiffman and Ralph, 2017). Nevertheless, additivity leads to some questionable predictions, especially for the initial Fl cross (Fraïsse et al., 2016). Second, while Fisher’s model is closely linked with quantitative genetics (Barton, 1989; Orr, 1998b; Rockman, 2012; Fisher, 1930, Ch. 2), a formal connection cannot be made without analytical approximations (Simon et al., 2018), and it is not clear how widely these approximations apply.

For these reasons, the current paper is in three parts. In Part I, we introduce the analytical predictions of Fisher’s model, focusing on the geometrical and biological meanings of its parameters. We also compare the approximations to individual-based simulations, including a wide range of assumptions about environmental change, population demography, and the population genetic regime. In Part II, we connect Fisher’s model to quantitative genetics, showing how the composite effects correspond neatly to the model parameters. We then express results for standard line crosses in different environments, unifying results from previous studies (Chevin et al., 2014; Hatfield and Schluter, 1999; Lynch, 1991; Rundle and Whitlock, 2001; Simon et al., 2018; Wright, 1922; Yamaguchi and Otto, 2019). Finally, in Part III, we introduce two extensions to the model, involving selectively induced associations between heterospecific alleles, and phenotypic dominance. These extensions address cases where the simplest model gives misleading or implausible predictions. We end by discussing some implications of our results for understanding the process of speciation.

## Model and Results

### 1 Fisher’s geometric model and hybridization

#### 1.1 Basics

We consider hybrids between two diploid parental lines Pl and P2, which differ by *d* substitutions. For simplicity, we will assume that the genetic variation within each parental line, is negligible compared to the divergence between them, although in principle, all analyses could be extended to include within-line variation (e.g. Roze and Blanckaert, 2014; Lynch and Walsh, 1998, Ch. 9).

Hybrids will contain some combination of alleles from the two parental lines. We characterize hybrid genotypes in terms of their heterozygosity, *p*_12_, and hybrid index, *h* (see Figure 1). The heterozygosity is the proportion of the *d* divergent sites where the hybrid carries one allele from each line; the hybrid index is the total proportion of the divergent alleles which come from line P2. As such, *h* ranges from 0 for a pure Pl genotype, to 1 for a pure P2 genotype. We will also use the notation *p*_l_ and *p*_2_ to refer to the proportion of divergent sites that are homozygous for alleles from Pl and P2, such that *p*_l_+*p*_2_+*p*_12_ = 1, and

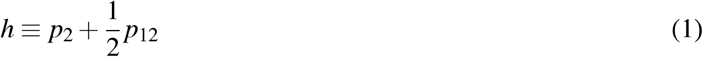

**Figure 1.**
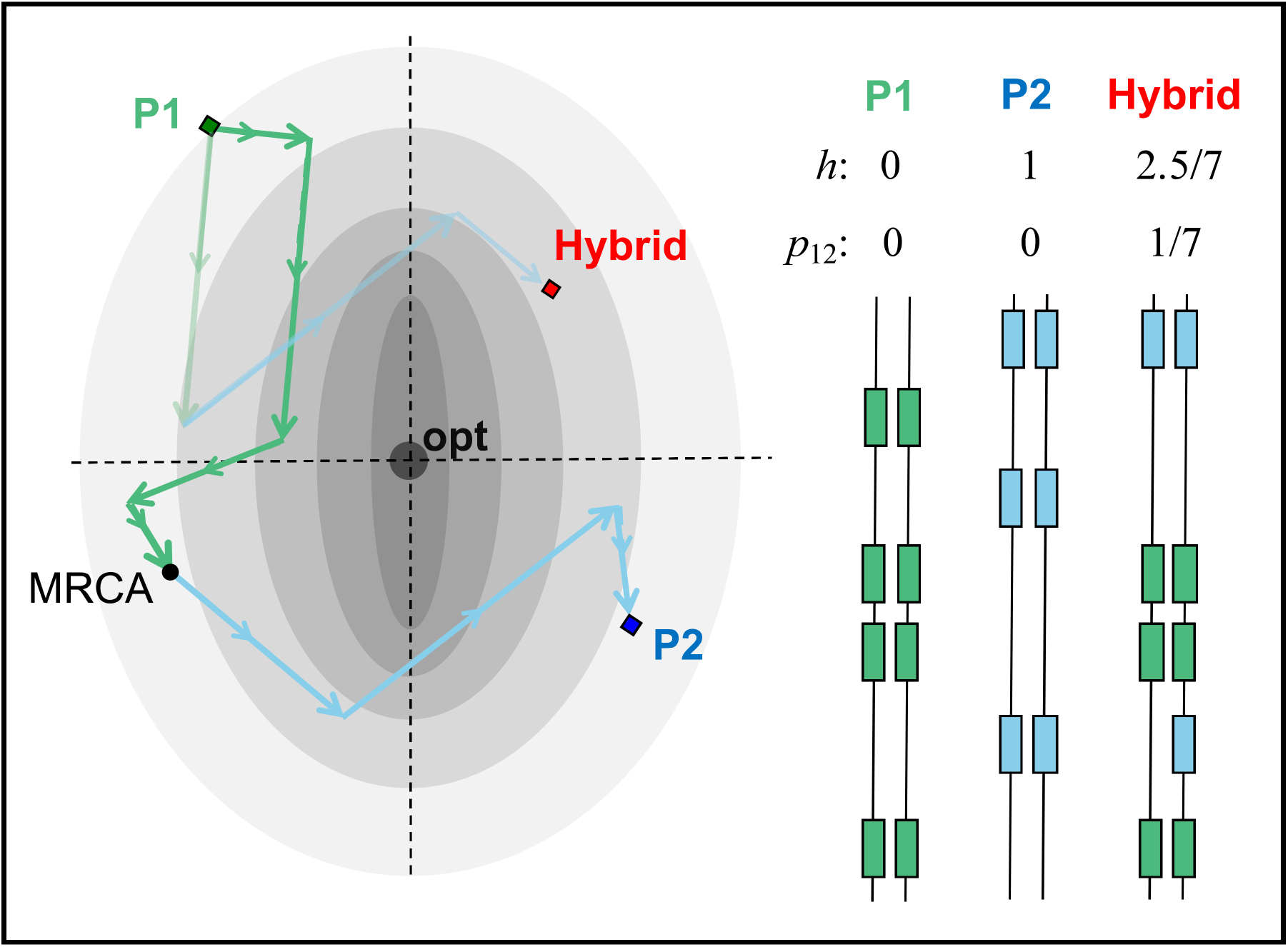
Fisher’s geometric model and hybridization. Each genotype is associated with the values of n quantitative traits (illustrated with *n* = 2), and so mutations and substitutions are vectors of change in this *n*-dimensional space. Fitness depends on the distance of the phenotype from an optimum. Shown are two parental lines, Pl and P2, which differ by *d* = 7 substitutions. The arrows represent the effects of each substitution in heterozygous and homozygous state. They are defined relative to the Pl genotype, regardless of whether the alleles are derived or ancestral (the *m*_*ij*_ in eq. 2 are the components of these vectors). Also shown is a hybrid genotype. In this hybrid, 1/7 of the divergent sites contains an allele from each line (*p*_12_ = 1/7), and two further P2 alleles are present as homozygotes (one ancestral and one derived), yielding a hybrid index of *h* = 2.5/7.

(Simon et al., 2018; Turelli and Orr, 2000). Our overall aim is to determine how the distribution of hybrid fitnesses depends on *h* and *p*_12_.

#### 1.2 Fisher’s model as a fitness landscape

Under Fisher’s geometric model, all genotypes are associated with the values of *n* continuously varying phenotypic traits. To begin with, we assume that the phenotypic effects of all *d* substitutions act additively on each trait. In this case, the value of trait *i* for any given hybrid genotype can be written as

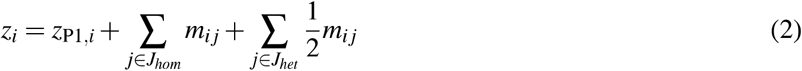

where *z*_P1,*i*_ denotes the value of trait *i* in parental line Pl, *J*_*hom*_ denotes the subset of *d* × *p*_2_ loci that are homozygous for the P2 allele, and *J*_*het*_ denotes the non-overlapping subset of *d* × *p*_12_ loci that are heterozygous. The *m*_*ij*_ describe the effects on trait *i* of introducing the P2 allele at locus *j* (whether derived or ancestral) into a Pl background. In the P2 genotype, by definition, all substitutions are present in homozygous state, and so its phenotype can be written as:

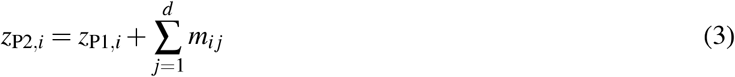

All of these quantities are illustrated in Figure 1. Next, we must decide how trait values determine fitness. We will assume that the fitness of a genotype depends on the distance of its phenotype from some optimal phenotype, as determined by the environment. We use the following weighted distance measure

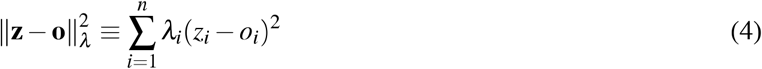

where **o** = (*o*_1_, *o*_2_, …, *o_n_*) is the vector of optimal values, and *λ*_*i*_ is the strength of selection on trait i in the current environment. (This metric is equivalent to the “breakdown score” of Turelli and Orr, 2000, and Simon et al., 2018.) Equation 4 implies that selection is independent on all traits. However, if correlated selection can be approximated by a multivariate Gaussian function, then, without loss of generality, we can always change the axes, and define n new traits, under independent selection (Martin and Lenormand, 2006a; Waxman and Welch, 2005). Furthermore, if the distribution of the fixed effects, the *m*_*ij*_, is also sufficiently close to multivariate normality, including arbitrary covariances, then we can choose new traits, under independent selection, which have unit variances and no covariances. As such, the *λ*_*i*_ capture both differences in selection, and differences in the typical sizes of factors fixed (Martin and Lenormand, 2006a; see also Appendix 1 for a full derivation).

Finally, fitness is a decreasing function of distance (Simon et al., 2018; Turelli and Moyle, 2007). This could take a form such as

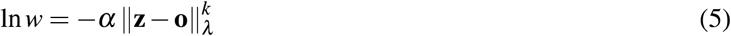

where *k* denotes the curvature of the fitness landscape, i.e. how quickly fitness declines with the distance from the optimum (Fraïsse et al., 2016; Fraïsse and Welch, 2019; Peck et al., 1997; Tenaillon et al., 2007). In the remaindernder of the paper, following common practice (Demuth and Wade, 2005; Turelli and Moyle, 2007; Turelli and Orr, 2000), we will not work with fitness directly. Instead, we follow (Simon et al., 2018), and use the following scaled squared distance (or scaled transformed fitness):

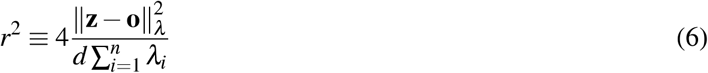

The denominator in eq. 6 can be understood in various ways. First, it is simply a convenient scaling to remove nuisance parameters, so that the results depend on a handful of simple geometric quantities. Second, when the parental lines are close to the optimum, applying the scaling is equivalent to dividing all fitness measures by the measures for some reference class of hybrid (Hill, 1982; Simon et al., 2018; Yamaguchi and Otto, 2019; Lynch and Walsh, 1998, Ch. 9). Third, the denominator can itself be written as a “reference distance”:

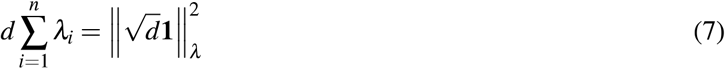

(where **1** is the unit vector). Equation 7 is the amount of phenotypic change that would be expected for a single population that wandered away from its ancestral state via an undirected random walk of *d* steps, where the step size is given by the *λ*_*i*_.

For brevity in what follows, “distance” will refer to scaled quantities such as eq. 6, and we return to the interpretation of the scaling below.

#### 1.3 The Brownian bridge approximation

Equations 2-3 suggest an approximation that can help us to understand hybrid fitnesses. In this section, we describe this approximation in geometric terms, and discuss its biological interpretation in the following section. If each hybrid genotype contains an effectively random sets of alleles, we can treat the *m*_*ij*_ as random variables that are statistically independent on each trait. This means that, if we consider a hybrid genotype with a given level of heterozygosity and hybridity, its expected distance from the optimum is

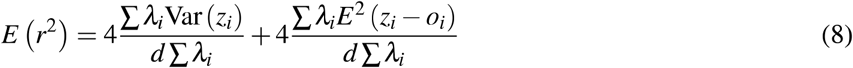

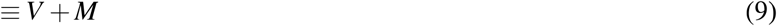

where the means and variances are defined over all hybrid genotypes with the same heterozygosity and hybrid index. To derive the quantities *V* and *M*, we will now assume that the *m*_*ij*_ on each trait are the increments of a Brownian bridge, i.e., a random walk constrained at each end by the parental phenotypes, and split into *d* steps (Revuz and Yor, 1999; Simon et al., 2018). This approximation relies on the fact that alleles are placed into hybrids in an effectively random combinations, but it does not require that the true process of divergence resembled a random walk. The approximation also depends on the additivity and normality of the *m*_*ij*_; but if the divergence, *d*, is sufficiently large, then the summations in eq. 2 lead to central-limit-type behaviour, and so to approximate normality, in a wider range of cases (Barton et al., 2017, section 3.2).

Simon et al. (2018) presented this Brownian bridge approximation, and in Appendix 1, we rederive the same result with less haste. For the term *V*, which captures the variation in hybrid phenotypes, we show that

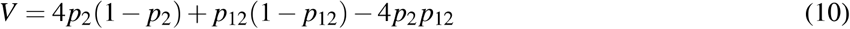

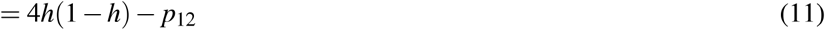

where the three terms in eq. 10, come from variation in the effects of homozygous P2 alleles, variation in the effects of heterozygous alleles, and a negative covariance term (because a given allele cannot appear in both homozygous and heterozygous state in the same genome). Note that the scaling of eq. 6 was chosen so that 1 ≥ *V* ≥ 0.

For the term *M*, we note that the expected hybrid phenotype will lie on the line connecting the parental phenotypes. *M* will therefore depend on the distance of the parental phenotypes from the optimum, and from each other. In particular, we find

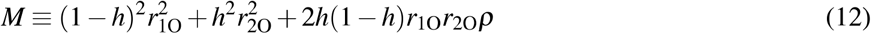

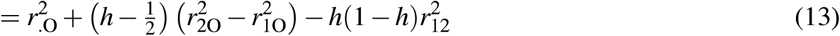

Equation 12 depends on three geometrical quantities, which are illustrated in Figure 2a. These quantities are the scaled squared distance to the optimum of two parental phenotypes:

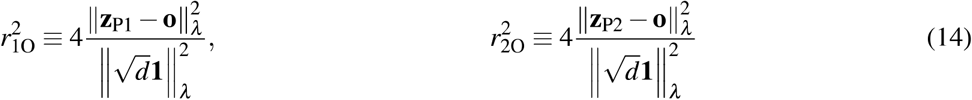

and *ρ*, which is the “cosine similarity” of the vectors connecting these phenotypes to the optimum. *ρ* can vary between 1, when these vectors point in the same direction, and −1, when they point in opposite directions (see also eq. 43). In equation 13, we use the cosine rule 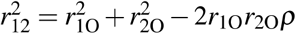, to give the same result in terms of the scaled distance between the parental phenotypes

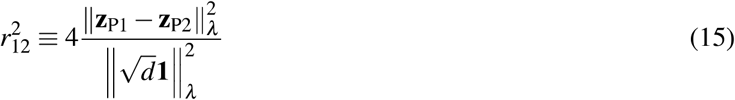

(Figure 2a) where we have also used the notation 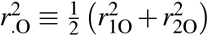 for the mean distance from the optimum of the parental phenotypes.

**Figure 2.**
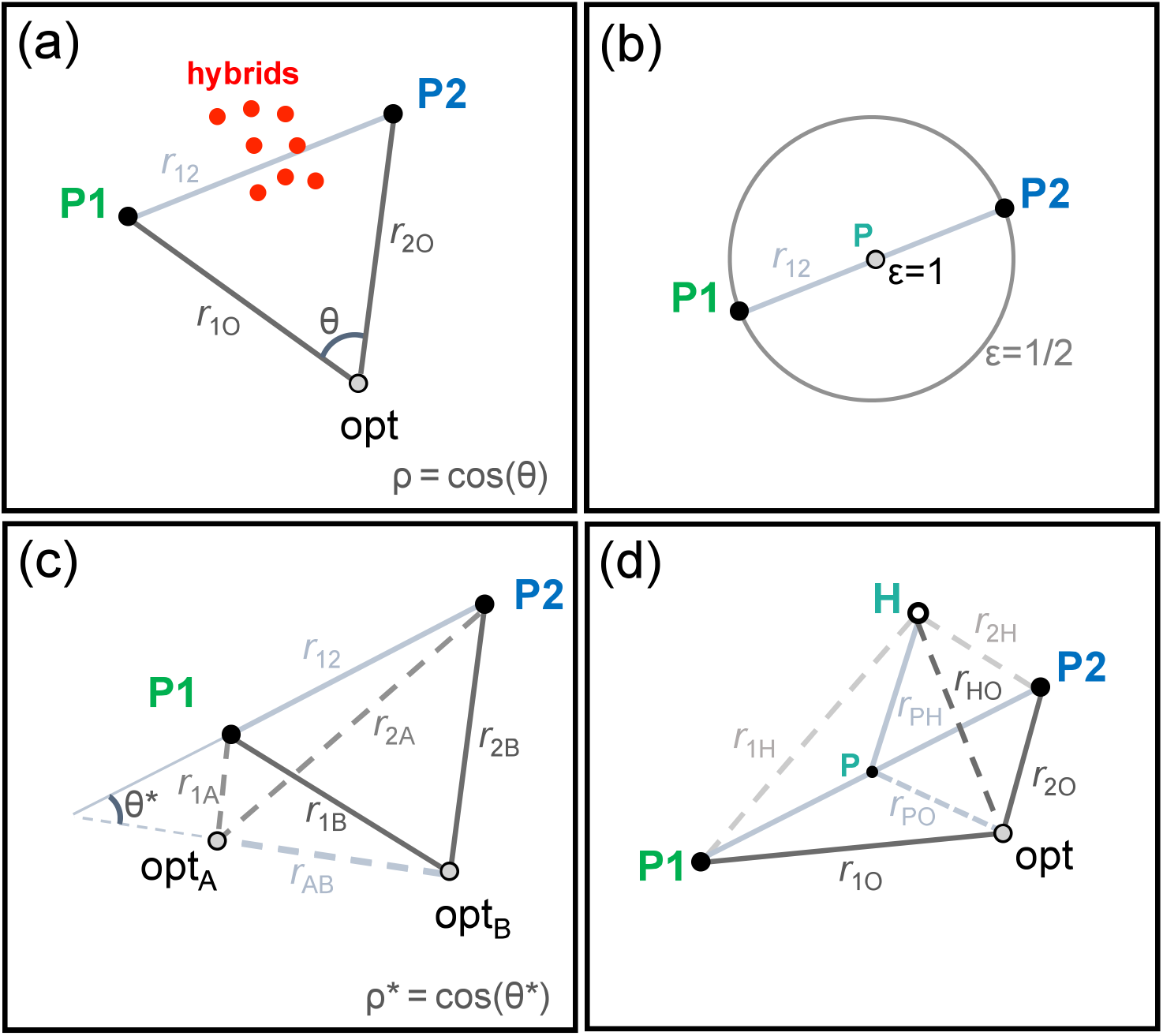
The predictions of Fisher’s geometric model for hybrids, depend on a small number of geometric quantities. These are defined in some multi-dimensional trait space, but are estimable, in principle. (a) With additive phenotypes, and a single environmental optimum, predictions depend on the scaled distance of the two parental phenotypes from the optimum (r_1O_ and r_2O_), and the angle between the vectors linking these phenotypes to the optimum. The cosine of this angle, *ρ*, measures the extent to which the parental populations are maladapted to the environment in similar ways. Results can also be given in terms of the scaled distance of the parental phenotypes from each other (r_12_), and this does not depend on the position of the optimum. (b) For balanced hybrids (with *h* = 1 /2), results depend solely on r_12_, and a quantity *ε*, which quantifies the potential of the current environment to generate hybrid advantage. The maximum value of *ε* = 1 is obtained with an intermediate optimum matching the midparental phenotype P, which corresponds to the average phenotype of balanced hybrids. Lower values of *ε* correspond to optima placed on circles of increasing diameter. (c) With two environments, A and B, characterized by different optima, one measure of local adaptation is *ρ*\*: the cosine similarity between the vectors linking the optima, and the parental phenotypes. When the two parental phenotypes are very close to the two optima, scaled hybrid fitness depends only on r_12_. (d) With variable phenotypic dominance, results depend on the phenotype of the global heterozygote, H, which is equivalent to the Fl cross under strictly biparental inheritance and expression. A consequence of variable dominance is that H may differ from the midparental phenotype, P. In the example shown, H is closer to the P2 phenotype than the Pl phenotype, this implies directional dominance, with Pl alleles being recessive on average.

#### 1.4 Biological interpretation

Let us now consider a group of hybrids that might vary in their values of *h* and *p*_12_. Combining results above, the expected distance of these hybrids from the optimum is

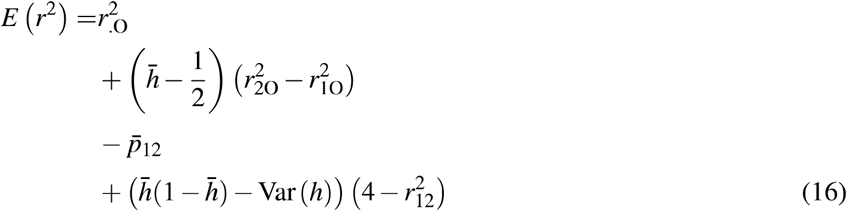

where 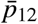 is the mean level of heterozygosity in hybrids of interest, and 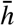 and Var(*h*) are the mean and variance of their hybrid indexes (Simon et al. 2018). All four of the terms in eq. 16 have a clear biological interpretation. First, and simplest, 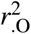 is the mean transformed fitness of the parental lines; it tells us that hybrids will be fitter, on average, if their parents are fitter, on average. The second term depends on the difference in the parental fitnesses: 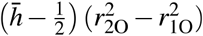; it tells us that hybrids will be fitter if they contain more alleles from the fitter parent. The third term, 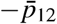, is an intrinsic benefit of heterozygosity; it states that, for any given value of *h*, hybrids are fitter when they are more heterozygous. This is a form of heterosis (Frankel, 1983). The effect is “intrinsic” because, unlike the two previous terms, it does not depend on the position of the environmental optimum. The final term captures the intrinsic effects of hybridity. Because it depends on 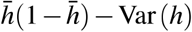, its contribution is smallest when hybrids are close to the parental types, either because 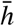 is close to 0 or 1, or because Var(*h*) is close to its maximal value: 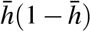. This effect is also “intrinsic”, because it depends on the distance 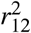, whose value is not affected by the position of the optimum (Figure 2a). The sign of this term changes with the size of 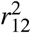. When 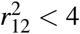, hybridity brings a net fitness cost. This reflects the breaking up of coadapted gene complexes in the parental lines (Lynch 1991; Simon et al., 2018; Wallace, 1991). When 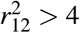, hybridity brings a net benefit. This reflects the potential benefits of transgressive variation in hybrids (Yakimowski and Rieseberg, 2014).

##### 1.4.1 Conditions for hybrid advantage

Results above imply that hybrids can sometimes be fitter than the parental lines. In particular, hybrids will be closer to the optimum, on average, than the average parental type, whenever 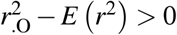.

To understand the conditions for hybrid advantage, it is simplest to consider balanced hybrids, with equal contributions from the two parental lines. With *h*= l/2, eq. 16, yields

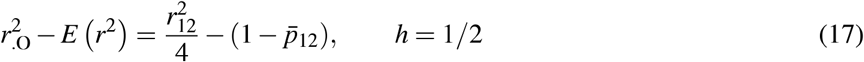

It follows that homozygous balanced hybrids (with *p*_12_ = 0), will enjoy a fitness advantage whenever 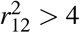. Fully heterozygous hybrids (with *p*_12_ = 1), will enjoy an advantage whenever 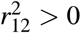, i.e. whenever the parents are phenotypically distinct. In general, the level of hybrid advantage increases with heterozygosity, and with 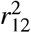, whatever the position of the optimum. The importance of environmental conditions is clearer from the relative hybrid advantage:

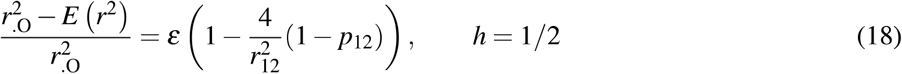

Here, a new quantity, 0 ≥ **ε** ≥ 1, describes the position of the optimum, and is defined by

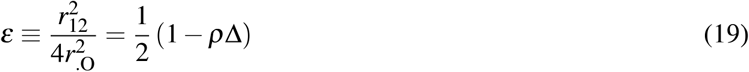

where *ρ* is defined above (Figure 2a), and 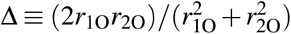, parameterises the difference in the distances of the two parents to the optimum. The quantity **ε** is illustrated in Figure 2b.

From eq. 18, the potential for hybrid advantage decomposes neatly into a property of the parental lines 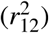, and a property of the optimum (**ε**). Advantage is maximized at **ε** = 1, when the parental lines are equally maladapted to current conditions (Δ = 1), and maladapted in opposite directions in phenotypic space (*ρ* = −1). In this case, the optimum coincides with the phenotype of the midparent (denoted Pin Figure 2b). This restates a result of Yamaguchi and Otto (2019), and also follows intuitively: intermediate hybrid phenotypes will best match intermediate environments (Moore, 1977).

##### 1.4.2 Hybrid fitness and the process of divergence

Results above show that the outcomes of hybridization depend strongly on the value of 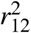. So what exactly does this quantity measure? By definition, 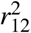 depends on the amount of phenotypic change that accrued during the genomic divergence between the parental lines; and this is scaled by the expectations under a random walk with the same distribution of effect sizes (eq. 15). For this reason, as shown in Figure 3, the value of 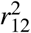 contains some information about the actual mode of divergence between the populations.

**Figure 3.**
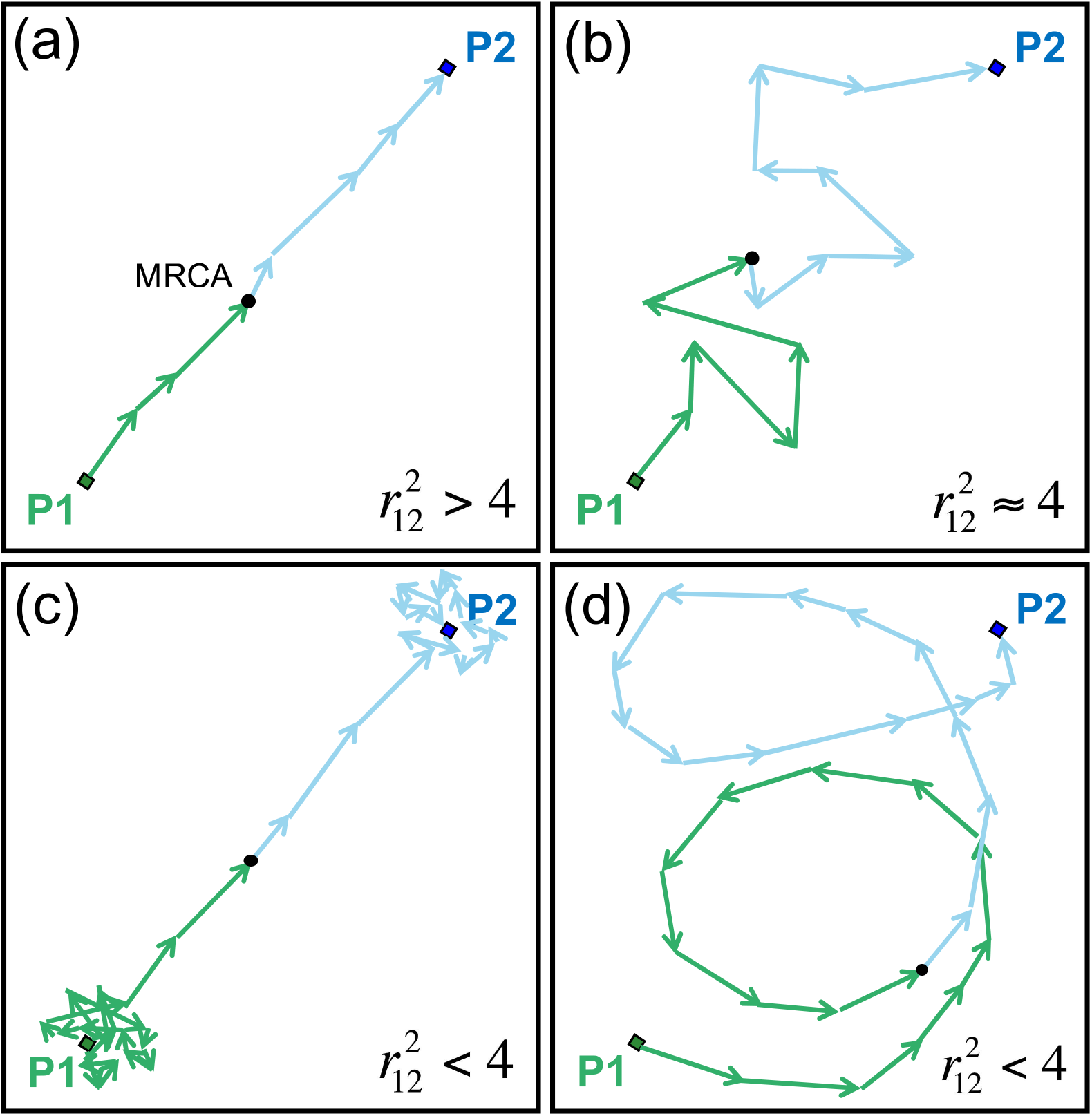
The scaled distance 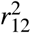 plays a key role in detennining patterns of hybrid fitness, and it can vary systematically with the mode of divergence between the parents. The variation depends on the chain of *d* substitutions that differentiate the parental lines, and compares their trajectory to a random walk with the same number of steps, and distribution of effect sizes. (a) When the substitutions form a more-or-less direct path between the parental phenotypes, the observed phenotypic difference will be greater than would be predicted under a comparable random walk; this implies that 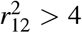. (b) when the true path of divergence really did resemble a random walk, then 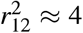 is expected. This might happen if stabilizing selection on the phenotype was ineffective, or if the optimum value wandered erratically. Systematically smaller values of 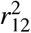 are predicted under two conditions. Either (c) genomic divergence continued, despite effective stabilizing selection on the phenotype, leading to “system drift”. Or (d) populations successfully tracked environmental optima, but without leading to a straight path of substitutions. If both lines are well-adapted under either of these last two scenarios, 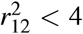 indicates coadaptation among the divergent alleles.

For example, if 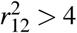, then the parents show more phenotypic divergence than expected under a random walk. This implies that the *d* substitutions, which connect the parental phenotypes, form a chain with relatively little meandering or changing of direction. Indeed it follows from the geometry that 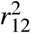 has a maximum at *4d*, when all of the substitutions cause identical changes in the same direction (see Appendix 2 for full details). Such a pattern is unlikely to arise without selection, and so an observation of 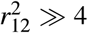 suggests that the parental lines diverged via positive selection, either acting in one population alone, or in both populations, but in opposite directions in phenotypic space. This is illustrated in Figure 3a.

Similarly, an observation of 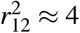 is expected if the parental phenotypes really did diverge by random-walk-like evolution (Figure 3b). One way this might occur is if parental lines fixed mutations regardless of their fitness costs, e.g. under severe inbreeding. It is notable that the intrinsic effects of hybridity vanish when 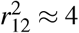 (eq. 16). This agrees with the empirical observation that heterozygosity, rather than hybridity, is the major determinant of fitness in crosses between inbred lines (Neal, 1935; Simon et al., 2018; Wright, 1922).

Finally, if 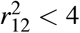, then populations have accrued less phenotypic divergence than would be expected under a random walk. This could occur in two quite distinct ways. First, stabilizing selection might maintain the phenotype at a (more-or-less) stationary optimum, while still allowing for divergence at the genomic level, perhaps by nearly neutral evolution (Barton, 1989; Hartl and Taubes, 1998). This process is closely related to “system drift” (Rosas et al., 2010; Schiffman and Ralph, 2017), and is illustrated in Figure 3c. Alternatively, divergence could involve adaptation to a moving optimum, but without leading to a straight path of substitutions connecting Pl and P2 (Mani and Clarke, 1990; Walsh and Lynch, 2018, Ch. 12). In the simplest case, this could arise if the two populations adapted, independently, to identical environmental change, because the chain of substitutions would then change direction as it passed through the common ancestor. A more complex example, involving an oscillating optimum, is illustrated in Figure 3d. In both cases, the result is a chain of substitutions whose start and end points are closer together than would be expected under a random walk, such that 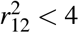.

Importantly, over large periods of time, at least one of these two processes - system drift, or complex environmental change - is very likely to occur. So at very large divergences, it becomes increasingly likely that 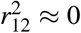 will hold, with the loss of any potential for hybrid advantage. In fact, it follows directly from eqs. 14–15 that all of the key distances shown in Figure 2a will tend to vanish at large divergences, and we have the limit:

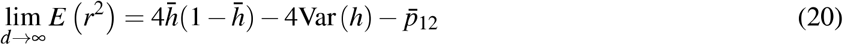

Biologically, eq. 20 implies that, as populations diverge genetically, both extrinsic fitness effects, and any intrinsic benefits of hybridity, will tend to become less and less important. The model predicts convergence to a characteristic pattern of intrinsic isolation between the parental lines, where a fixed cost of hybridity is mitigated by a fixed benefit of heterozygosity.

#### 1.5 Individual-based simulations

To illustrate the points above, and test the robustness of our approximations, we used individual-based simulations of Fisher’s model. In particular, we simulated the divergence between pairs of populations, followed by hybridization in the ancestral environment. We simulated under 2^7^ = 128 different combinations of population-genetic parameters (Table 1). These parameters were chosen to encompass a range of conditions, including varying levels of standing variation, recombination, distributions of mutant effects, and other properties of the fitness landscape. Justifications of our parameter choices, and full details of all simulations, are given in Appendix 3.

**Table 1:**
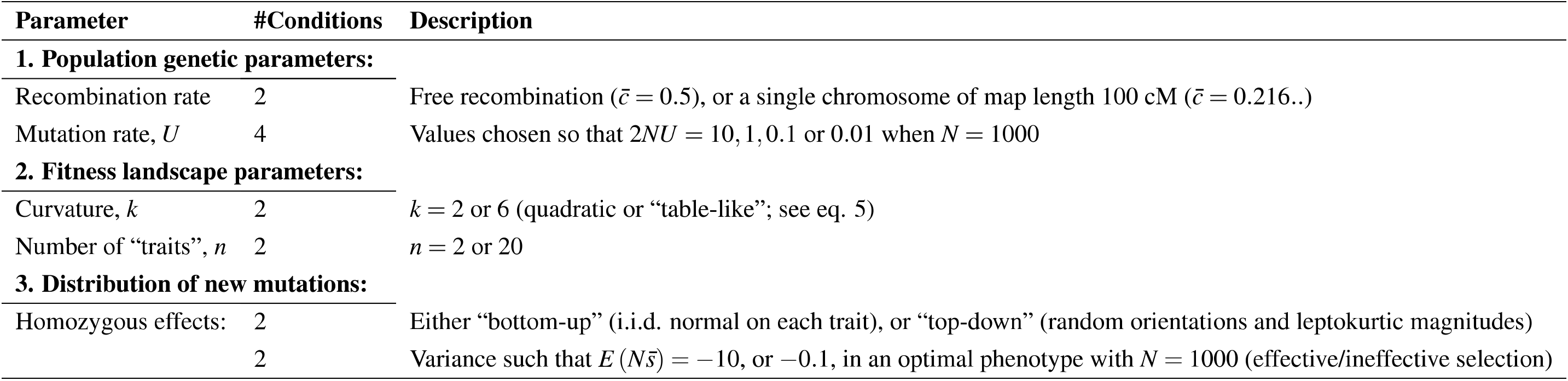
Parameter values used in the simulations.

For each of the 128 parameter combinations, we simulated 15 distinct scenarios of population demography, and environmental change. The purpose of the scenarios was to generate predictably different patterns of genomic divergence (Figure 3), and predictably different patterns of hybrid fitness in the ancestral environment (Figure 2b).

Figure 4 describes these divergence scenarios, and plots the realized values of the key quantities 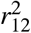 and **ε** (see eqs. 15, 18 and 19). First, let us consider the results for scenarios 1-4. In these scenarios, the divergence between the populations involved local adaptation, with optima that moved in opposite directions, with respect to the ancestral habitat (see top right-hand panel). As Figure 4 shows, the adaptive divergence in these simulations led to some very high values of 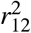, conducive to hybrid advantage. The 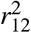 values are highest when the divergence took place in the face of ongoing gene flow (scenarios 1-2). This is because the gene flow tended to keep the overall genomic divergence low (Yeaman and Whitlock, 2011), such that paths of substitutions resemble Figure 3a more than Figure 3c. The 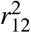 values are also higher when the optima moved in single jump (scenarios 1-3), than when it moved back and forth via smooth cycles (scenario 4). This reflects the meandering in the chains of substitutions when environmental conditions fluctuate (compare Figure 3d to Figure 3a). For all four scenarios, values of *ε* were close to their maximal value of 1, implying that the optimum in the ancestral habitat - which was exactly intermediate between the two new optima - was ideally placed to maximize hybrid advantage (Figure 2b). There was also some variation in *ε*, either because the cycling optima returned to their ancestral state (scenario 4), or because high levels of maladaptive gene flow moved the parental populations away from their local optima (scenario 1).

**Figure 4.**
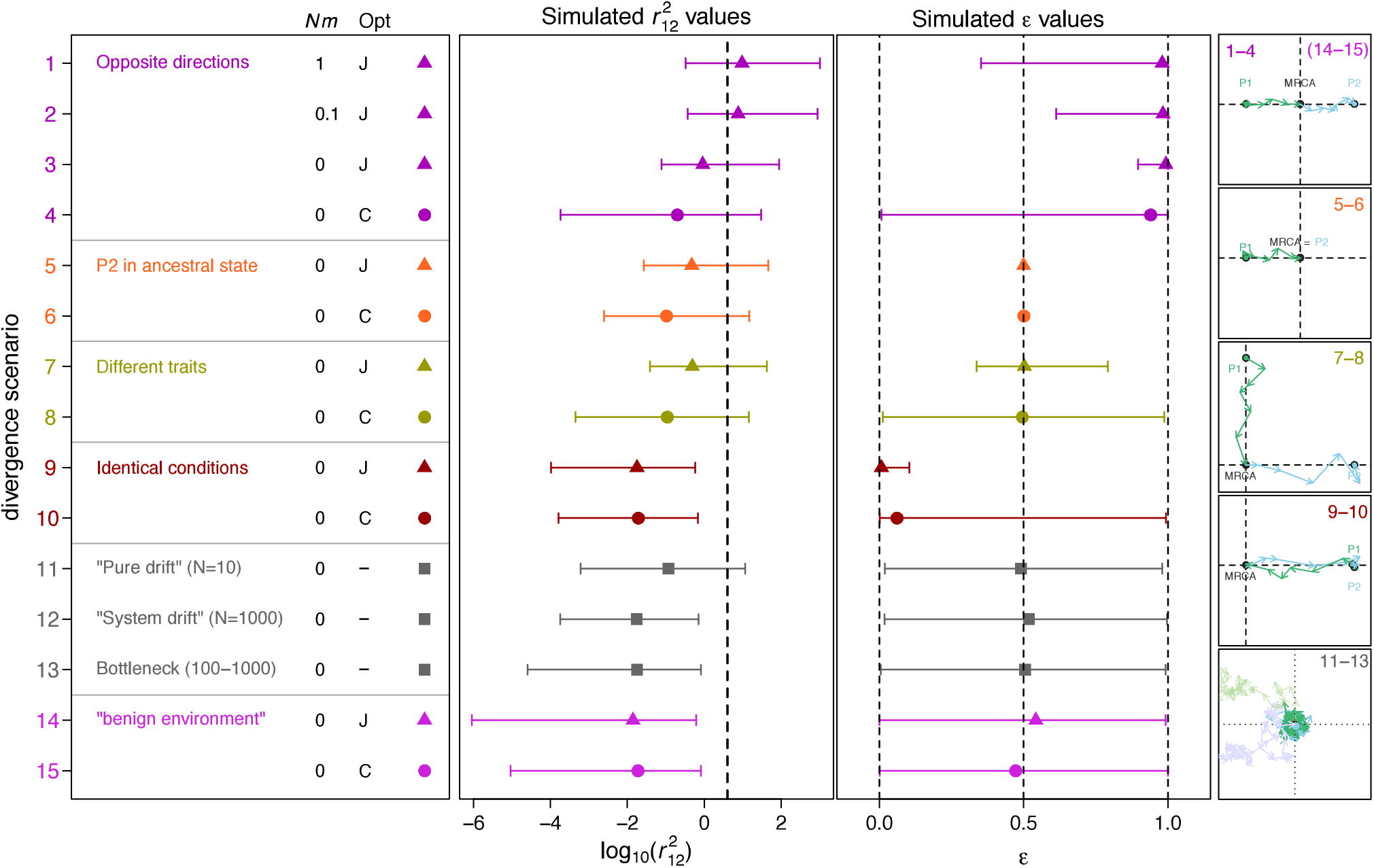
Individual-based simulations of population divergence under Fisher’s geometric model. We simulated 15 distinct divergence scenarios, summarized in the left-hand panel. These scenarios differed in the rate of gene flow during divergence(captured in the expected number of migrants, *Nm*), and the pattern of environmental change. In particular, optima moved in a single abrupt jump(“J” and triangular points), in smooth repeating cycles(“C” and circular points), or remainderned stationary throughout the simulation(“-” and square points). Patterns of environmental change also varied between the scenarios as indicated by colour, and illustrated in the right-hand panels. The two central columns show the realized values of two important determinants of hybrid fitness; with points and intervals showing the median and complete range of 128 simulations runs, each with the different set of population genetic parameters (see Table 1). The second column shows 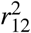: the scaled difference between the parental phenotypes (Figure 2b), whose value can reflect the process of divergence(Figure 3). The vertical dotted line shows the value 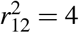, above which homozygous hybrids have an expected fitness advantage over their parents. The third panel shows *ε*; this describes the position of the optimum in a given habitat, such that expected hybrid fitness increases with *ε* (Figure 2b). Values shown here are for the ancestral habitat, which matches the common ancestral phenotype of the diverging populations. For allopatric simulations(scenarios 3-15) runs terminated after one of the populations had fixed 500 substitutions, such that d ≤ 1000. For these runs, simulated values of 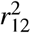 can be compared to the possible maximum log_10_(4d) ≈ 3.6. For parapatric simulations(scenarios 1-2), runs terminated after 50 */U* generations, and mutations were treated as “fixed” if allele frequencies differed by more than 50% between the demes; 11 runs, for which d < 10, were excluded. Full details of the simulations are found in Appendix 3.

Scenarios 5-6 resemble scenarios 3-4, except that population Pl evolved, while population P2 remainderned in their common ancestral state (see second right-hand panel). As expected, this had little effect on 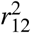 values, but reduced e from around 1 to exactly 1/2. This is because the ancestral optimum no longer maximized hybrid advantage. In the ancestral habitat, hybrids were often fitter than parental line Pl, but never fitter than the optimal P2 (Figure 2b). Very similar results were obtained for scenarios 7-8. Here, populations adapted to moving optima on different traits (see third right-hand panel). In these cases, the value of **ε** ≈ 1/2 reflected the fact that hybrids could gain a fitness advantage over both parental lines, but less so than in an intermediate habitat (Figure 2b).

In scenarios 9-10, results were qualitatively different. Here, the populations adapted, independently, to identical environmental change (see fourth right-hand panel). As a result, the parental phenotypes were very close to each other (such that 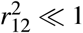), and the ancestral optimum was poorly placed to generate hybrid advantage (**ε** ≈ 0).

The next group of scenarios, 11-13, involved stable environmental conditions, so that all of the divergence accumulated via genetic drift in allopatry (bottom right-hand panel). Here, values of **ε** varied greatly, reflecting the fact that parental populations deviated from their shared optimum in independent, random directions. High values of 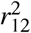 were obtained only in one case: when the population size was very small (scenario 11). In this case, selection was often ineffective, at least initially, so that the population phenotypes wandered away from their optimal values (see Figure 3b, and lighter lines in the lower right-hand panel of Figure 4). By contrast, for scenarios 12-13, parental phenotypes remainderned close to their shared optimum at all times (see darker lines in lower right-hand panel of Figure 4), and this kept values of 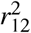 consistently low.

Finally, for scenarios 14-15, we reanalysed the simulations from scenarios 3-4, but scored hybrids in a “benign environment” (Hatfield and Schluter, 1999). This means that fitness was affected only by the *n* − 1 phenotypic traits that had not been subject to diversifying selection. As such, the fitness effects in hybrids reflect the pleiotropic effects of adaptive substitutions (Thompson, 2019), and results closely resemble those under “system drift”.

Figure 4 confirms that there are consistent differences between hybridization scenarios, which reflect both the history of divergence (captured in 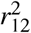) and the current environmental conditions (captured in **ε**). Nevertheless, shown in Figure 5a–b, the Brownian bridge approximation applied well to all of the simulations reported above. Figure 5a shows results for hybrids that were formed by randomly assembling combinations of homozygous alleles from the parental lines, in different proportions (i.e., with different values of *h*). Results show that 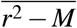 is well predicted by eq. 11 for all simulations. In many of these simulations, the distribution of fixed effects was highly non-normal (see Figure Sl for details), but the Brownian bridge approximation applied nonetheless, as shown in Figure 5b illustrates another feature common to all of the simulations: *M* tended to vanish if the genomic divergence grew large. With large *d*, the outcome of hybridization was always predicted by eq. 20, regardless of the divergence scenario, or the current position of the optimum.

**Figure 5.**
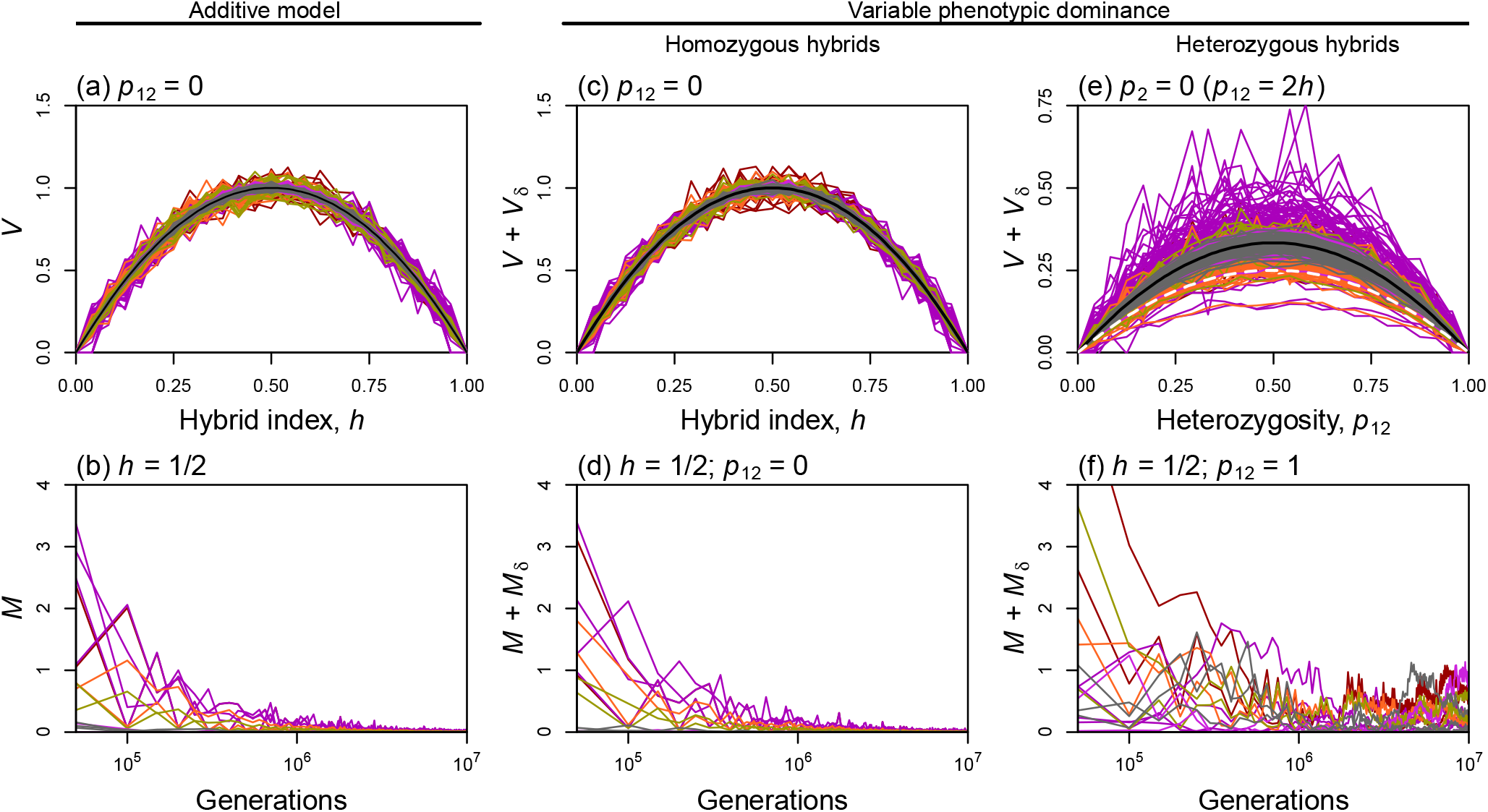
The Brownian bridge approximation (eqs. 8–13) compared to individual-based simulations. (a) Predictions for *V* (eq. 11), apply well to all divergence scenarios and population genetic parameter regimes. The lines plotted are the mean of *r*^2^ − *M*, across 10,000 homozygous hybrids, created from *hd* P2 alleles, and (1 − *h*)*d* Pl alleles, chosen at random. Results are shown for all of the simulations reported in Figure 4, and use the same colour scheme. (b) For all divergence scenarios, *M* tends to decline over time, as long as genetic divergence increases (eq. 20). Results are shown for balanced hybrids (*h* = 1/2), with an illustrative set of parameters (*k* = 6, *n* = 2, *NU=* 1, *Ns* = −0.1, bottom-up mutations, and free recombination). Similar results were obtained for all parameter regimes, except for a subset of simulations of divergence in parapatry, where divergence levels reached a quasi-equilibrium value, instead of increasing steadily. (c)-(f) Adding variable phenotypic dominance to the model yields qualitatively different results for heterozygous hybrids (eqs. 28–30). (c) The Brownian bridge approximation continues to apply to homozygous hybrids, but (e) a second Brownian bridge applies to heterozygous alleles placed in a Pl background. The white dotted line shows the prediction of *p*_12_(l − *p*_12_) that applies under additivity; while the black line shows the prediction of (1 + 4*v*)*p*_12_(l − *p*_12_) with *v* = 1/12. (f) *M* +*M_δ_* declines over time for homozygous hybrids, but (f) not for heterozygous hybrids. For balanced hybrids, with *h* = 1/2, this reflects the difference between the midparent (with *p*_12_ = 0, 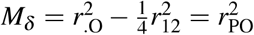) and the global heterozygote (with 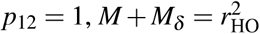). In most cases, the parental phenotypes (and therefore the midparent) are directly subject to selection, while the global heterozygote is not. Results imply that - with phenotypic dominance - Fl fitness can be low at high divergences.

### 2 Fisher’s model and the quantitative genetics of line crosses

In this second part of the paper, we show how the distances in phenotypic space, which govern the fitness of hybrids, relate to measurable quantities. To do this, we will consider controlled crosses, including the initial Fl (Pl×P2), the F2 (random mating among the Fl), and backcrosses of the Fl to the parental lines. These crosses vary in the key quantities 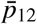, 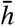 and Var(*h*), and the last of these also depends on the levels of segregation and recombination (Lynch and Walsh, 1998, Ch. 9). In particular, if 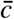 is the mean rate of recombination among pairs of loci, then 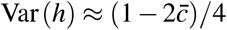 among Fl gametes (Zeng et al.,1990; Lynch and Walsh, 1998, Ch. 9); and so, with random union of gametes, Var(*h*) will be half of this value for the F2, and a quarter of this value for backcrosses. Furthermore, 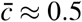 for many species (Lynch and Walsh, 1998, Ch. 9), and in those cases, Var(*h*) can often be neglected.

#### 2.1 The composite effects under Fisher’s model

Let us begin by following Hill (1982; see also (Lynch, 1991; Lynch and Walsh, 1998, Ch. 9)), and writing the expected value of a trait in a hybrid as

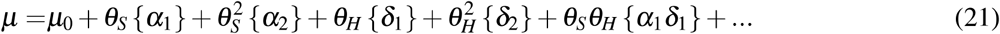

where 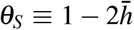 and 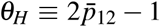. The curly brackets contain the “composite effects”, which are defined in Table 2 (and noting that, in this standard notation, {*α*_1_*δ*_1_} describes an interaction term, and not a product). Equation 21 includes only pairwise effects, but the model might also be expanded to include higher-order terms.

**Table 2:**
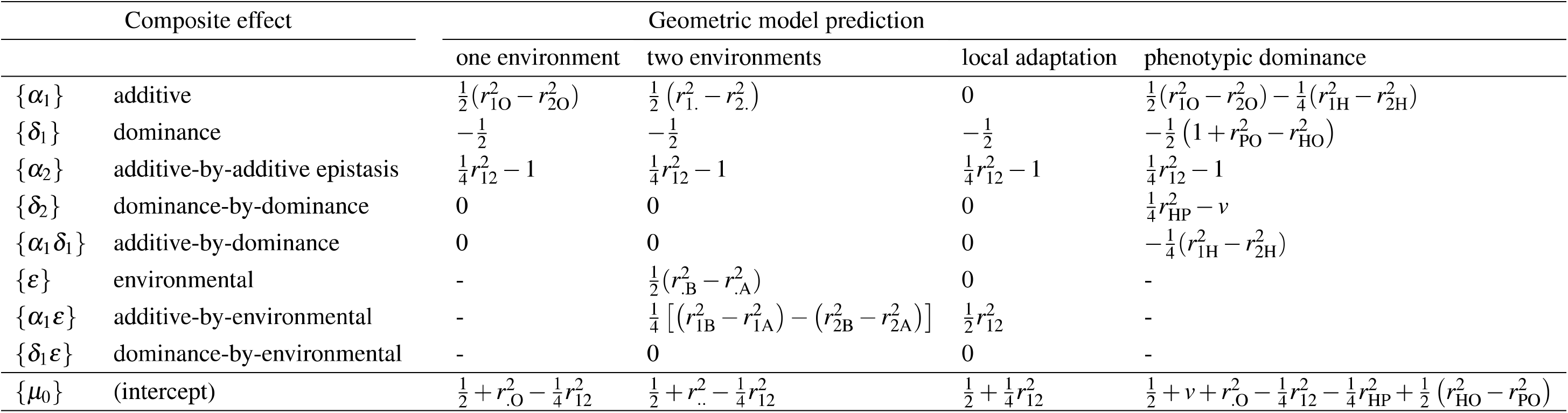
Composite effects under Fisher’s geometric model.

If we neglect Var(*h*), then both eqs. 16 and 21 are polynomials in 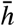 and 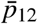, and so we can apply eq. 21 to the transformed hybrid fitness, setting 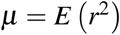, and solve for the composite effects. The results, found in Table 2 (column “single environment”), show that Fisher’s model predicts three non-zero composite effects. Their values reflect the biological distinctions discussed above. In particular, the additive effect, 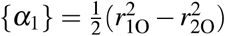, captures the benefits of carrying alleles from the fitter parent, while the dominance effect, 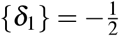, captures the intrinsic benefits of heterozygosity. The pairwise epistatic effect, 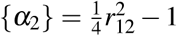, balances the intrinsic costs and benefits of hybridity.

Rundle and Whitlock (2001) presented a useful extension of eq. 21 for traits scored in two environments. Introducing an indicator variable, *I*, which is O for individuals scored in “environment X’, and 1 for individuals scored in “environment B”, and defining *θ*_*E*_ ≡ 2*I* − 1, their model contains the additional terms:

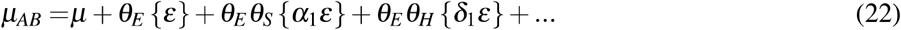

Fisher’s geometric model is trivially extended in the same way, by adding a second environment, with a distinct optimum. This is illustrated in Figure 2c. Again, we can solve for the composite effects, and these are shown in Table 2 (column “two environments”). Results show that adding a second environment leaves the dominance and epistatic effects unchanged, confirming that they represent the intrinsic effects of heterozygosity and hybridity. Of the remainderning quantities, the additive effect, {*α*_1_}, is now averaged across environments (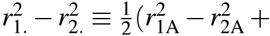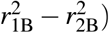), while the main environmental effect, {**ε**}, is simply the difference in fitness between environments, averaged across the parental lines 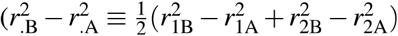.

Finally, the additive-by-environment interaction is

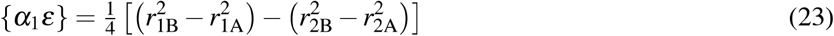

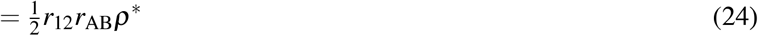

Here, −1 ≤ *ρ*\* ≤ 1 is the cosine similarity of the vector linking the parental phenotypes, and the vector linking the two optima (Figure 2c): {*α*_1_**ε**} will be large when the difference between the phenotypes of Pl and P2 resembles the difference between optima A and B. Indeed, the predicted values of {*α*_1_}, {**ε**} and {*α*_1_**ε**} are equivalent to the quantities described by (Blanquart et al., 2013) for measuring the extent of local adaptation (see their eqs. 1–2), but applied to fitness values that have been transformed and scaled (eq. 6). As shown in Appendix 4, the same framework is also extendable to hybrids that are formed in patchy ecotones, or in habitats that are ecologically intermediate between A and B.

#### 2.2 Local adaptation and ecological isolation

While {*α*_1_**ε**} is a possible measure of local adaptation, it does not describe the extent of ecological isolation between the parental lines. For example, {*α*_1_**ε**} might be very high, even if Pl were fitter than P2 in both habitats (Blanquart et al., 2013; Kawecki and Ebert, 2004). However, we do have a measure of isolation in an important special case. If the two parental lines are well adapted to different local optima, then 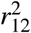 describes the distance between these optima, as well as the distance between the parental phenotypes. As a consequence, results for hybrids depend on 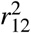 alone, and this is shown in Table 2 (column “local adaptation”).

With local adaptation of this kind, large values of 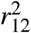 imply pure ecological isolation (where P2 and Pl are kept distinct solely by environment-dependent selection against their divergent alleles), while small values imply pure intrinsic isolation (where hybrids are less fit than the parents in both habitats). By changing the value of 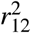 we can interpolate between these two extremes, and this is shown in Supplementary Figure S2. With local adaptation, 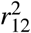 is both a measure of the amount of meandering in the chain of fixed differences, and a measure of the relative strengths of ecological isolation versus intrinsic isolation.

To illustrate this, Figure 6 shows results from a single simulation run. In this run, parental populations adapted in allopatry, to distinct optima. The adaptation involved fixing ~50 substitutions, and after this, the populations continued to diverge via system drift. We consider hybridization between these populations in five distinct environments (Figure 6a), tracking both the composite effects over the complete course of the divergence (Figure 6b–e), and the fitness of crosses generated by standard amphimixis, at two time points (Figure 6f–o).

**Figure 6.**
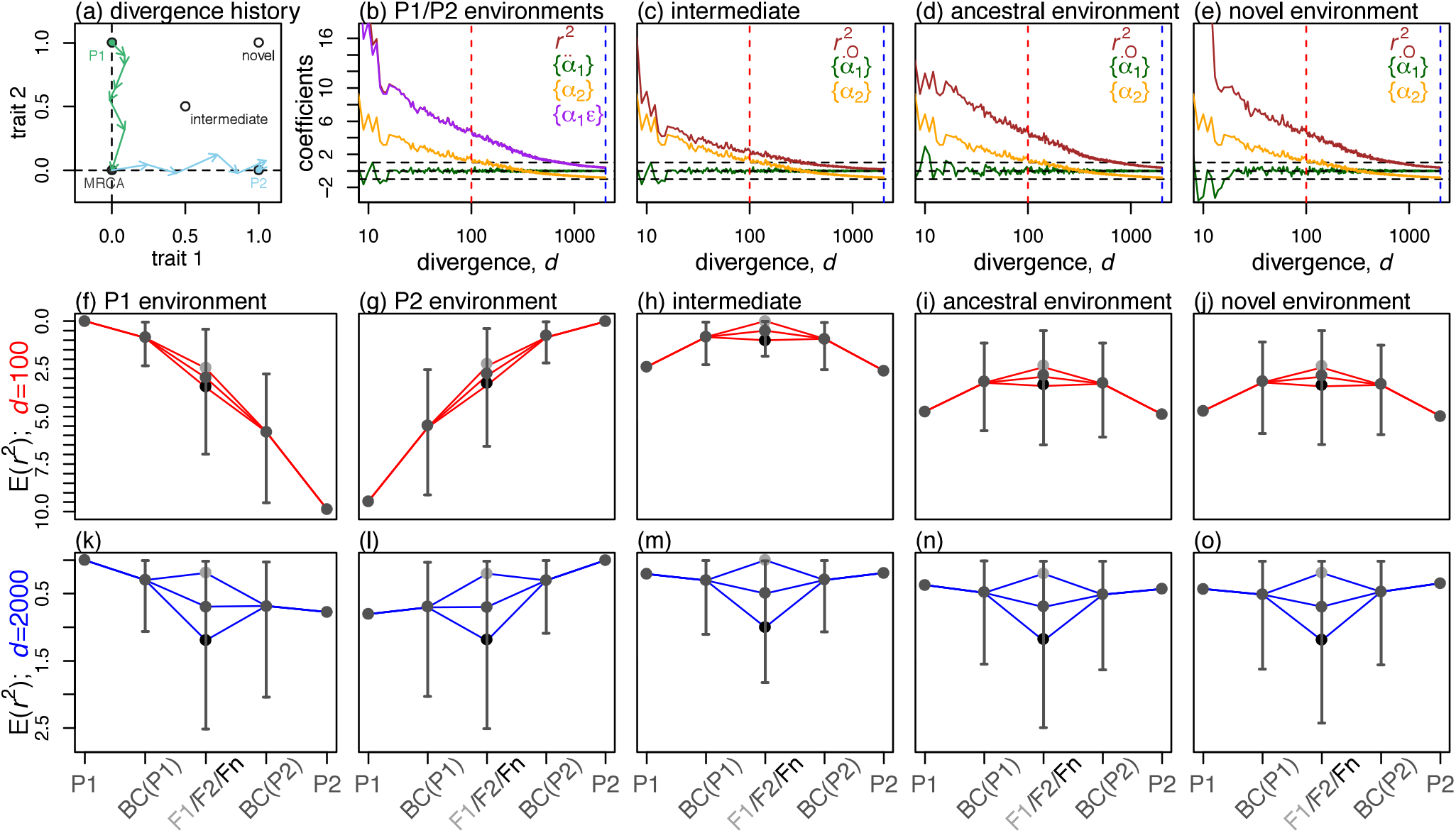
Hybridization with local adaptation under Fisher’s geometrical model. Panel (a) shows a cartoon of the divergence process that was simulated, with two populations adapting to abruptly shifting optima, and then continuing to accumulate divergence in allopatry. In the common ancestral population, the initial values of all traits were set to zero, but the optimal value of one of the traits was set to one, choosing a different trait for each population (i.e., “scenario 7” from Figure 4). Panels b-e show the change, with increasing divergence, in the composite effects (Table 2), as measured with respect to both parental environments (panel b), or in other single environments (panels c-e). The remainderning panels show results for simulated hybrids, plotting *r*^2^ on a reversed axis, such that fitter genotypes are higher up on the plots. Points show simulated values for the three fixed genotypes (the parental lines, Pl and P2, and the Fl = Pl × P2). Points with error bars show mean and 95% quantiles for 10,000 recombinant hybrids, from the reciprocal backcrosses: BC(Pl) = Fl × Pl and BC(P2) = Fl × P2; and the F2 = Fl × Fl. The dark point at the centre of each plot shows results for homozygous hybrids, derived as from automictic selfing among Fl gametes. As such the three central crosses (Fl, F2 and “Fn”, are all balanced hybrids, with 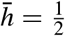, but have maximally different heterozygosities, 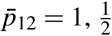 and 0). All points are compared to analytical predictions (blue and red lines). These use eq. 16, with the measured values of *r*_1O_ and *r*_2O_ and the assumption that *ρ* = −1 (for the “intermediate” environment) or *ρ* = 0 (all other cases). We note that these predictions relied on our knowing the scaling factor (eq. 7) which can be measured directly from simulated data, but not from real-world data (see Appendix 1 and Discussion). Finally, hybrids were scored in the early stages of divergence (*d* = 100; red lines, panels f-j), where the signal of local adaptation is strong, and at later times (*d* = 2000; blue lines, panels k-o), where the composite effects tend to decline, with convergence towards a characteristic pattern of intrinsic isolation (eq. 20). Divergence was simulated with the following parameters: *N* = 1000, 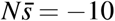, *2NU* = 1, *n* = 2, *k* = 2, free recombination, and “bottom-up” mutations (see Table 1).

As is clear from Figure 6b–e, several of the composite effects change in similar ways, reflecting their common dependence on 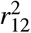. In the earliest stages of divergence, 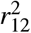 values were high, reflecting the adaptation of the parental lines to distant, and distinct optima. After 100 substitutions, this was manifest in the clear signatures of ecological isolation in the parental habitats (Figure 6f–g). In the environment to which Pl is adapted, hybrids tended to be fitter when they carry more Pl alleles, and vice versa (Rundle and Whitlock, 2001). For the same reason, in an ecologically intermediate habitat, there was a clear signal of bounded hybrid advantage (Figure 6c). Hybrids tended to have the favoured intermediate phenotype (Moore, 1977; Yamaguchi and Otto, 2019). Hybrid advantage, at a lower level, also occurred in the ancestral habitat (Figure 6i); but this had nothing to do with the habitat being ancestral, and the same patterns were observed in an entirely novel habitat, characterized by similar values of 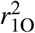, 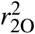 and 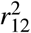 (Figure 6j).

As the genomic divergence increased, 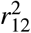 decreased, and so all of these differences between environments were greatly reduced (Figure 6b–e). In the example shown, after ~2000 substitutions, hybrid fitnesses were already converging towards the characteristic pattern of intrinsic isolation (Figure 6k–o), with the fixed cost of hybridity, {*α*_2_} ≈ −1, and the fixed benefit of heterozygosity 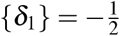 (eq. 20).

The results shown in Figure 6 assume free recombination between all loci, but comparable results apply with limited recombination (Supplementary Figure S3). Convergence to the characteristic pattern of intrinsic isolation occurs whatever the mode of divergence (Supplementary Figures S4-S5).

### 3 Two extensions

In this third and final part of the paper, we highlight two ways in which the model gives misleading or implausible predictions, and show how these limitations might be overcome.

#### 3.1 Later crosses, and the Bulmer effect

Results above rely heavily on heterospecific alleles appearing in random combinations, but with later generation hybrids, selection on the earlier generation hybrids can induce non-random associations between alleles in their gametes. This can increase the fitness of their offspring, without changing allele frequencies in the population as a whole (Bulmer, 1971; Walsh and Lynch, 2018, Ch. 16). For example, with random union of gametes, the distributions of *h* and *p*_12_ will often remaindern unchanged between the F2 and F3 generations, and so equations 16 and 21 make the same predictions for both. However, selection on the F2 parents can upset this expectation.

To see this, let us consider the case of free recombination 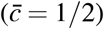, and optimal parental lines (*r*_.O_ = 0). With these assumptions, the variance in trait values (the *z*_*i*_) among F3 offspring is

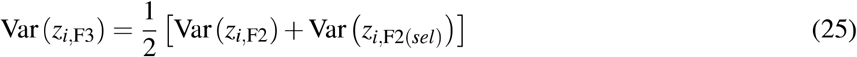

where *z*_*i*,F2_ and *z*_*i*,F2(*sel*)_ are the trait values for the total F2 population, and for the subset of selected parents (see Walsh and Lynch, 2018, Ch. 16, assuming complete heritability). In the extreme case, if only optimal F2 reproduce, then Var (*z*_*i*,F2(*sel*)_) = 0 and 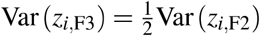. Because the trait variance halves, we have the general bounds

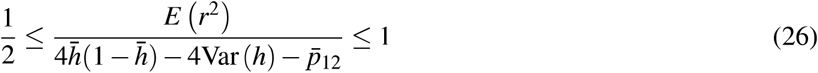

where the upper bound comes from eq. 20, and the lower bound applies if only optimal individuals reproduce. Later crosses will fall somewhere between these bounds, depending on the strength of selection acting on the earlier generation hybrids. This is illustrated in Figure 7a–b. We chose two simulation runs where populations diverged despite a fixed optimum, via system drift. If we generated an F3 cross using a random selection of F2 parents, then the upper bound of eq. 26 applies well (see blue lines in Figure 7). If we selected parents with a probability proportional to their fitness (eq. 5), the same results continued to apply for later crosses, but only when the fitness function was quadratic (*k* = 2), so that there was very little inter-individual variation in fitness (Figure 7a). When we imposed a “table-like” fitness function (*k* = 6; Fraïsse et al., 2016), equivalent to strong truncation selection, and implying high variation in parental fitness, then results were closer to the lower bound of eq. 26 (see red lines in Figure 7b). These patterns continued unchanged for other late generation crosses, including the F4 and F5, and also applied to repeated backcrosses to the Pl line (Figure 7a–b).

**Figure 7.**
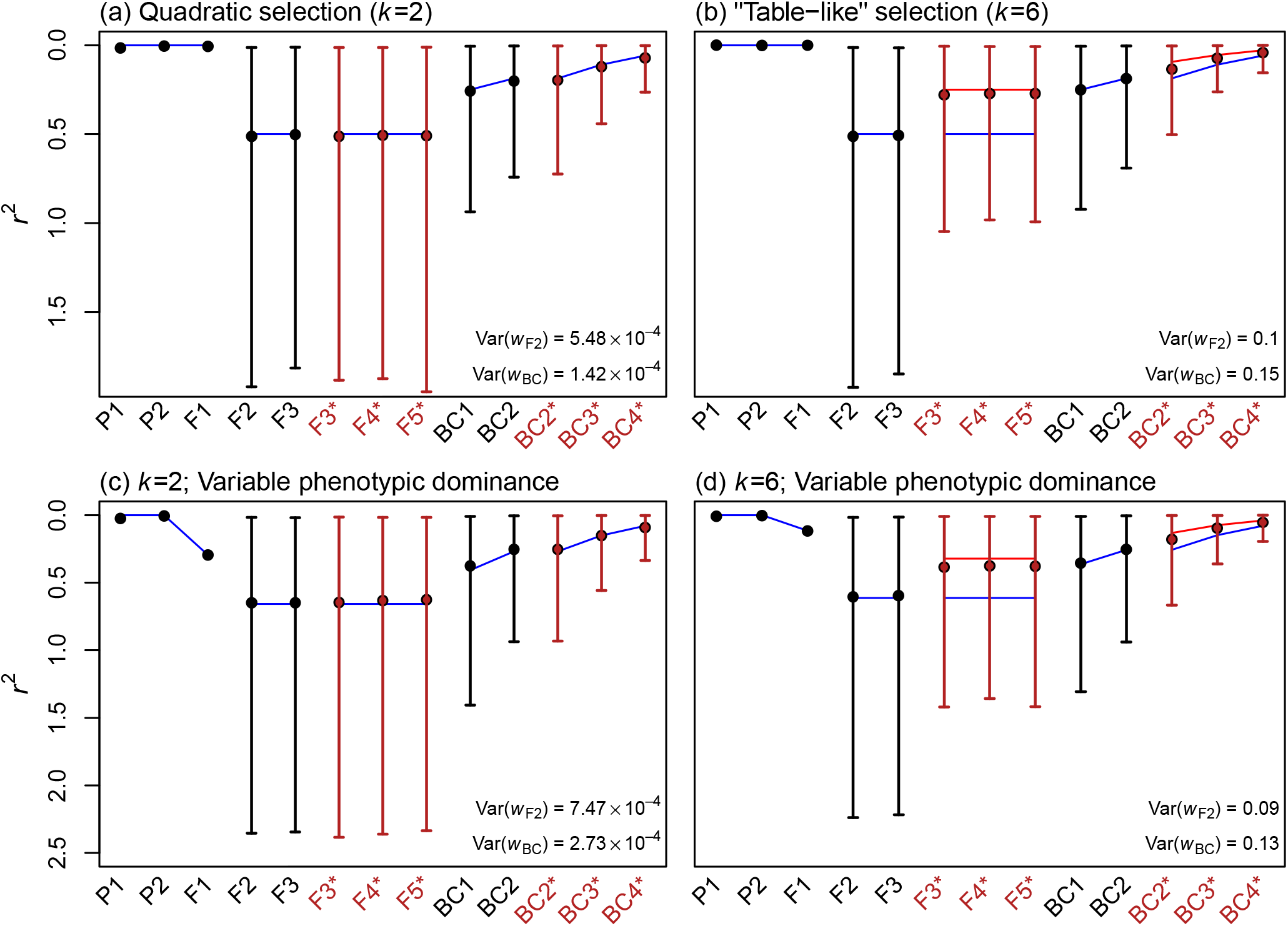
Hybrid fitness for later-generation crosses, showing the effects of selection acting on the earlier hybrids. Plots show the squared distance from the optimum, *r*^2^, on a reverse axis, so that higher points are fitter. After the initial Fl cross, we simulated either random union of gametes among the hybrids (F2-F5), or repeated backcrossing to parental line Pl (BC1-BC4). For the later crosses, we chose parents either wholly at random (black points and lines), or with a probability proportional to their fitness (asterisks and red points and lines). In each case, results for 10,000 simulated hybrids (mean and 95% quantiles), are compared to analytical predictions from eq. 16 (blue lines), assuming optimal parental phenotypes (*r*_.O_ = 0). This is equivalent to the upper bound of eq. 26. When the variance in parental fitness was low (panel (a)), selection had little effect. When variance was high (panel (b)), results for the later generations approach predictions for extreme truncation selection, such that only optimal parents reproduce (red lines; lower bound of eq 26). Panels (c)-(d) show equivalent results, when populations were simulated with variable phenotypic dominance. The clearest consequence is that the Fl are suboptimal, even when the parental lines are optimal (blue and red lines used eqs. 29 and 30 with the observed 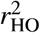, and *v* = 1/12, from the variance of a uniform distribution). All panels used simulations with the following parameters: *N* = 1000, 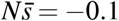, *2NU* = l, *n* = 2, free recombination, “bottom-up” mutations, stationary optima matching the ancestral state; hybrids were formed as soon as one of the diverging populations had fixed 1000 substitutions.

#### 3.2 Phenotypic dominance

The most implausible prediction of Fisher’s model is embodied in eq. 20. This equation predicts that fully heterozygous hybrids will always be as fit as their parents, because, when *p*_12_ = 1 and *h* = 1/2, the benefits of heterozygosity exactly cancel the costs of hybridity (Barton, 2001; Fraïsse et al., 2016; Schiffman and Ralph, 2017).

This prediction is worrying because - with strictly biparental inheritance and expression - the initial Fl cross will be fully heterozygous. While many intrinsically isolated species do produce fit Fl (Fraïsse et al., 2016; Price and Bouvier, 2002; Wallace, 1991), Fl fitness tends to decline as the parents become very genetically divergent, even in environments where both parents are well adapted (Bateson, 1978; Edmands, 2002; Endler, 1977; Fraïsse et al., 2016; Price and Bouvier, 2002; Waser, 1993). The model also struggles to explain a second widespread pattern in the Fl: when some loci have uniparental inheritance or expression, the reciprocal Fl often have very different fitnesses, even if the parents are both well adapted (Bolnick and Near, 2005; Bouchemousse et al., 2016; Brandvain et al., 2014; Escobar et al., 2008; Fraïsse et al., 2016; Sato et al., 2014; Turelli and Moyle, 2007). Fisher’s model can only account for these asymmetries if the global heterozygote is suboptimal (see Fraïsse et al., 2016 for details).

In this section, we show that these features of Fisher’s model are improved by adding phenotypic dominance (Manna et al., 2011). To see this, let us replace eq. 2 with

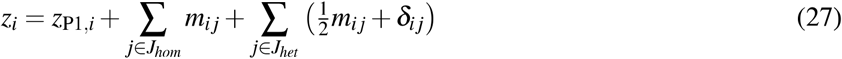

where *δ*_*ij*_ is the deviation from semi-dominance on trait *i*, caused by introducing the P2 allele at locus *j* in heterozygous form. We now assume that the *δ*_*ij*_ can be treated as the increments of a new, and independent Brownian bridge, linking the midparental value of trait *i*, to the trait value of the global heterozygote (see the phenotypes labelled P and H in Figure 2d). In Appendix 1, we show that this assumption leads to

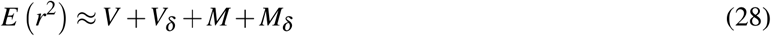

where *V* and *M* are the additive results (eqs. 10–13), and *V*_*δ*_ and *M*_*δ*_ are the new contributions from variable dominance. In Appendix 1, we show that these new contributions are:

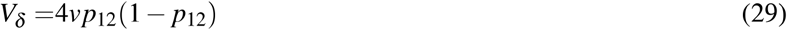

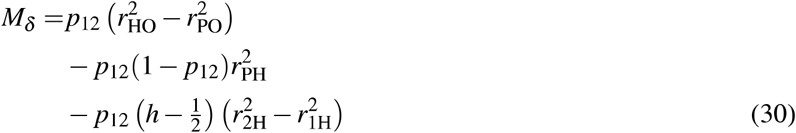

In equation 29, a new parameter, *v*, describes the scaled variance of the *δ*_*ij*_. Equation 30 depends on a number of scaled distances, and these are illustrated in Figure 2d. The corresponding changes in the composite effects are listed in Table 2 (column “phenotypic dominance”). Table 2 shows that phenotypic dominance adds two new composite effects: {*δ*_2_} and {**α**_1_*δ*_1_}, and alters the value of a third, {*δ*_1_}, so that it is no longer a constant.

Equations 29 and 30 are both proportional to *p*_12_ and so they alter predictions only for heterozygous hybrids. The predictions are altered in two ways. First, a non-zero value of {**α**_1_*δ*_1_} (which corresponds to the third term in eq. 30), now allows for “directional dominance”. For example, in Figure 2d, the global heterozygote, H, is much closer to the P2 phenotype than to the Pl phenotype (*r*_2H_ < *r*_1H_), which implies that P2 alleles are dominant on average. T his sort of asymmetry allows Fisher’s model to account for “dominance drive” in hybrid zones, where alleles can spread due to their dominance relations alone (Barton, 1992; Mallet and Barton, 1989). Second, by allowing the phenotype of the global heterozygote, H, to differ from the midparental phenotype, P, phenotypic dominance alters the effects of heterozygosity in hybrids. While these effects are intrinsic and beneficial under the additive model, with phenotypic dominance the effects can vary over time and space. Most importantly, heterozygosity will tend to become deleterious at large divergences. This is because the global heterozygote - unlike the parental genotypes - may never be exposed to natural selection, and cannot, in any case, breed true. As such, its phenotype can continue to wander away from the optimum as divergence increases, even if effective stabilizing selection acts on the parental phenotypes. The result is that the distance 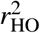, unlike 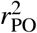, has no tendency to vanish as divergence increases.

To test the predictions of eqs. 28–30, we repeated our complete set of simulations (Figure 4; Table 1), but now including variable phenotypic dominance. To do this, the heterozygous effect of each new mutation on each trait, was generated by multiplying its homozygous effect by a uniformly-distributed random number. Results are summarised in Figure 5c–f, and in more detail in Supplementary Figure S6.

Figure 5c corresponds to Figure 5a, and shows that earlier results continue to hold for homozygous hybrids. Figure 5e shows equivalent results for heterozygous hybrids, i.e., random assemblies of P2 alleles, added in heterozygous state to a Pl background. In this case, because *h* = *p*_12_/2, we have the prediction *V* + *V*_*δ*_ = (1 + 4*v*)*p*_12_(l − *p*_12_). In most cases, results agree with this prediction, after setting *v* = 1/12, which matches the variance of the uniform distribution, used to generate the heterozygous effects of new mutations (see the black line in Figure 5e). However, the parameter *v* applies to fixed effects, and not to new mutations. As such, there are clear differences between the values of *v* realised in different parameter regimes and divergence scenarios, although usually heterozygous hybrids are less fit than would be predicted under an additive model, with *v* = 0 (see the dotted white line in Figure 5e; and Figure S6 for more details).

Results relevant to the Fl are shown in Figure 5d and 5f. These panels confirm that when divergence has increased, and the parental distances have declined towards zero (Figure 5d), the phenotype of the global heterozygote can remaindern a non-negligible distance from the optimum (Figure 5f). This implies that optimal but divergent parents will generate an unfit Fl. This is confirmed in Figure 7c–d. Here, results closely match the additive results shown in Figure 7a–b, except for the Fl, which is noticeably less fit. Similar results for a complete set of crosses are shown in supplementary Figure S7.

## Discussion

Using fitness landscapes based on Fisher’s geometric model, we have developed analytical predictions for the fitness of hybrids between divergent lines. These predictions apply to a wide range of evolutionary and ecological scenarios (e.g. Figure 4), and involve a small number of quantities with an intuitive biological meaning. For example, with the simplest model (additive phenotypes, and a single optimum), results depend solely on three distances: 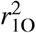, 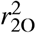 and 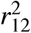 (Figure 2a), and so we can classify scenarios according to the values of these distances.

Next, we drew a formal connection between Fisher’s model, and the quantitative genetics of line crosses (Demuth and Wade, 2005; Hill, 1982; Lynch, 1991; Rundle and Whitlock, 2001; Yamaguchi and Otto, 2019). We showed that the key distances are closely related to the composite effects, and so can be estimated from cross data. This claim comes with an important caveat. The results in Table 2 apply not to raw fitness values, but to values that have been suitably transformed and scaled; in our notation, they apply to *r*^2^ and not to *w* (see eqs. 5–6). Data transforms are an inherent part of quantitative genetics (Lynch and Walsh, 1998, Ch. 11), but there is also the need to estimate the “reference distance”, which is the scaling factor in eq. 6. This extra parameter is relatively easy to estimate from a diverse collection of hybrids (see Simon et al., 2018), but not from a limited number of controlled crosses (Yamaguchi and Otto, 2019). We ducked this issue in Figure 6, by estimating eq. 7 directly from the simulated fixed effects (see Appendix 1). This is a real limitation, but there are many special cases where the distances can be estimated from fitness values that are transformed but unscaled (i.e., from the numerator of eq. 6). For example, with two locally-adapted populations (Table 2), the key distance 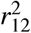 can be estimated from a ratio of composite effects:

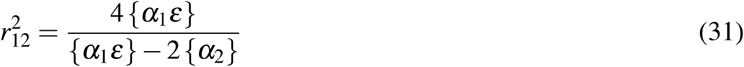

so that the scaling factor cancels. Simon et al. (2018) give other, similar examples.

In a third and final part of this work, we explored two extensions to Fisher’s model. The first extension involved disequilibria between heterospecific alleles, generated by selection on early generation hybrids (as opposed to selection during the divergence). These effects are weak in some parameter regimes (Figure 7a), but observations of strong incompatibilities, involving small genomic regions, suggests that they might be important in nature (Barton, 2001; Fraïsse et al., 2016; Coyne and Orr, 2004, Ch. 8). Accordingly, we provided a simple result, which applies with strong, truncation-like selection (eq. 26; Figure 7b). However, this approach will be difficult to apply to entire hybrid swarms, when some, but not all hybrids have strong selectively-induced disequilibria (Allendorf et al., 2001; Jiggins and Mallet, 2000; Simon et al., 2018; Vemesi et al., 2003). Even greater challenges will arise when selection changes allele frequencies (Walsh and Lynch, 2018, Ch. 16 and 24). In both cases, the distributions of *h* and *p*_12_ will not be sufficient to predict hybrid fitnesses.

In the second extension, we incorporated phenotypic dominance into Fisher’s model. Our simple treatment ignored a known feature of fitness dominance: the differences in the typical dominance relations of large- and small-effect changes (Fraïsse et al., 2016; Manna et al., 2011; Wright, 1929). Nevertheless, our treatment removed the most implausible prediction of the additive model, concerning the high fitness of the Fl. By reducing the beneficial effects of heterozygosity, the introduction of dominance allows for low fitness Fl between highly divergent, but equally fit parental lines (Fraïsse et al., 2016; Figure 7c–d). This extension also shows how Fisher’s model can incorporate other modelling approaches as special cases (Simon et al., 2018). For example, when the parental lines have high fitness, eq. 28 is identical to eq. A37 of Simon et al. (2018), which was derived from a model of Dobzhansky-Muller incompatibilities, with variable dominance relations (Torelli and Orr, 2000).

### The process of divergence and the outcome of hybridisation

Because it is a well-studied model of evolutionary divergence, Fisher’s model is especially useful for investigating the connections between the divergence process and the outcome of hybridisation.

One set of connections has been explored extensively in previous work. Compared to drift, positive selection will lead to divergence that is more rapid and more resistant to the swamping effects of gene flow, and tend to fix effects that are larger and more variable in size (Débarre et al., 2015; Dittmar et al., 2016; Griswold, 2006; Matuszewski et al., 2014, 2015; Orr, 1998b; Rockman, 2012; Thompson et al., 2019; Yamaguchi and Otto, 2019; Yeaman and Whitlock, 2011). Larger changes will often have a greater influence on hybrid fitness (Chevin et al., 2014; Fraïsse et al., 2016; Yamaguchi and Otto, 2019). Together, these facts will tend to implicate natural selection, rather than drift, in any hybrid problems that appear early in the divergence process (Jiggins and Mallet, 2000; Yamaguchi and Otto, 2019; Coyne and Orr, 2004, Ch. 11). In the results presented here, these effects of selection are all incorporated into the reference distance (eqs. 6–7), with selection tending to lead to larger values of *d* and larger values of the *λ*_*i*_.

The present work has focused on a different set of connections between divergence and hybridisation, and these are captured by the scaled distance 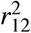. This distance can be called “intrinsic”, because it is a property of the parental lines, which does not depend on the current position of the optimum. For this reason, 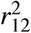 describes the possible outcomes of hybridisation in a variety of environmental conditions (Figure 6, Figure S2). For example, when parental lines are well adapted to different habitats, a high value of 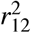 implies that the isolation between the lines will be purely ecological. In an intermediate habitat, or wherever the parents are poorly adapted, lines with a high 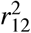 are more likely to generate hybrid advantage beyond the Fl. In this way, 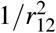 provides a natural measure of “coadaptation” among the parental alleles (Wallace, 1991). This is also clear from the fact that 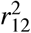 determines {*α*_2_}, the additive-by-additive epistatic effect (Table 2; Lynch, 1991).

As well as describing the outcomes of hybridisation, 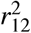 contains some information about the mode of divergence between the parental lines. This information is not about epistasis: the value of {*α*_2_} tells us nothing at all about the role of epistatic genetic variance during divergence (Barton, 2017; Demuth and Wade, 2005; Lynch, 1991; Welch, 2004). Instead, we have shown that 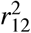 measures the consistency in the “directions” of the divergent alleles in trait space, and compares this consistency to expectations under a random walk. As such, high values of 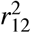 imply that substitutions had effects in the same direction more often than expected (Figure 3). This definition shows that 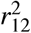 is closely connected to standard tests for natural selection on quantitative traits, such as the QTL sign test (Orr, 1998a), or the Qst-Fst comparison (Spitze, 1993; Whitlock and Guillaume, 2009). Indeed, we have confirmed that adaptive divergence, especially in parapatry, is most likely to lead to high values of 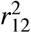 (Figure 4).

Together, these results clarify what hybrid fitness can and cannot tell us about the mode of parental divergence. On one hand, some patterns of hybrid fitness - those associated with high 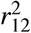 - are reliable indicators of selectively driven divergence, and especially of local adaptation maintained in the face of gene flow. On the other hand, patterns associated with low 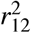 can arise in a variety of ways, including via adaptive divergence, especially in allopatry (e.g. Figure 3d). (These limitations are closely related to the low power of the QTL sign test: Rice and Townsend, 2012; Walsh and Lynch, 2018). Furthermore, unless there is substantial gene flow, any signature of selection will be transient. Over time, the model predicts convergence to an identical pattern of intrinsic reproductive isolation, whatever the mode of divergence (Figures 3–5).

## Acknowledgements

We are very grateful to Jean-Baptiste Grodwohl, Sally Otto, Aylwyn Scally and Alexis Simon. Some of the simulations used the Montpellier Bioinformatics Biodiversity platform supported by the LabEx CeMEB, an ANR “Investissements d’avenir” program (ANR-10-LABX-04-01). EDS and HS are funded by the Wellcome Trust programme in Mathematical Genomics and Medicine (codes PFHZ/157 and PCGG.GAAB respectively). HS also acknowledges financial support from the MEME programme in Evolutionary Biology.

## Author contributions

JW, NB, and HS designed the project. JW, EDS, and HS performed the analysis. HS, DR, and JW wrote the simulation code and ran the simulations. JW, HS, and EDS wrote the manuscript, with contributions from all authors.

