## Appendices for "The geometry and genetics of hybridization"

### Appendix 1:

In this Appendix, we derive the Brownian bridge approximation for Fisher's geometric model (eqs. 8, 12). Equivalent results were also presented by Simon et al. (2018), but using a different method.

##### A1.1 Removing correlations in the fitness function and fixed factors

After this section, we will assume that the fixed effects on each trait are uncorrelated, and will also assume uncorrelated selection on each trait, as in eq. 4. Before deriving the main results, we will first show that they apply more generally, as long as the distribution of fixed effects is sufficiently close to multi-variate normal, potentially involving correlations among traits, and as long as the distance metric is a multi-variate Gaussian, potentially involving correlated selection. This is achieved by transforming the phenotypic space. This "simultaneous diagonalisation", allows us to work with a far simpler class of models, without any loss of generality. Our treatment is very similar to that of (Martin and Lenormand, 2006a), and follows most notation from (Mathai and Provost, 1992).

We start with a vector $X$ of phenotypic effects in an $n$-dimensional space, which follows a multivariate normal distribution, that is,

$$X \sim N(\mu, \Sigma) \tag{32}$$

We also have an arbitrary quadratic form

$$Q(X) = X^T A X \tag{33}$$

where $A$ is positive definite. In the model, $Q(X)$ functions as a general formula for the distance to the optimum, which, without loss of generality, is assumed to be at the origin. In particular, as in Martin and Lenormand (2006a), the diagonal entries of $A$ are the selective effects of each trait, and off-diagonal entries correspond to selective interactions between traits. The positive definiteness of $A$ means that we cannot attain a negative value for $S(X)$, so that fitness is maximized at the optimum.

Now define

$$Y = \Sigma^{-1/2}(X - \mu) \tag{34}$$

so that $Y \sim N(0, I)$. Then diagonalize the matrix $\Sigma^{1/2} A \Sigma^{1/2}$ as

$$P^T \Sigma^{1/2} A \Sigma^{1/2} P = \Lambda \tag{35}$$

where $P$ is an orthogonal matrix so that $P^T P = I$ and $\Lambda$ is a diagonal matrix of eigenvalues of $\Sigma A$. Now let

$$U = P^T Y \tag{36}$$

so that $Y = PU$ and $U \sim N(0, I)$. Now we have

$$Q(X) = X^T A X = (Y + \Sigma^{-1/2}\mu)^T \Sigma^{1/2} A \Sigma^{1/2} (Y + \Sigma^{-1/2}\mu) = (U + b)^T \Lambda (U + b) \tag{37}$$

where $b = (P^T \Sigma^{-1/2} \mu)^T$. Lastly, defining $Z = U + b$, we have

$$Q(X) = X^T A X = Z^T \Lambda Z = \sum_{j=1}^{n} \lambda_j Z_j^2 \tag{38}$$

where $Z \sim N(b, I)$ is our transformed phenotypic vector with no correlation between traits, $\Lambda$ is our transformed, diagonal selection matrix. This procedure leads to the distance measure of eq. 4, and explains our use of isotropic selection in the simulations. Of course, in practice, we are unlikely to know the exact distribution of the phenotypic effects, and even from simulations, we can only measure their sample mean and covariance. This means that the normalising constant in eq. 6 has to be estimated in other ways (see Discussion).

##### A1.2 The Brownian bridge approximation

Having defined the independent traits, starting with eqs. 2-3, we now assume that random collections of heterospecific alleles describe a Brownian bridge on each trait. We do this by treating the homozygous effects of the P2 alleles (i.e., $m_{ij}$), as increments in the bridge, linking the trait values of P1 and P2. We note here that the factor 4 in eq. 6 appears for consistency with Simon et al. (2018), who considered a bridge applying to heterozygous alleles, and ending at the midparent; $E\left(r^2\right) \equiv f$ in their notation. From the standard properties of Brownian bridges (Revuz and Yor, 1999), we have:

$$E\left(m_{ij}\right) = \frac{z_{\mathrm{P2},i} - z_{\mathrm{P1},i}}{d} \tag{39}$$

$$\mathrm{Cov}\left(m_{ij}, m_{ab}\right) = \begin{cases} 0, & i \neq a \\ 1 - 1/d, & i = a, j = b \\ -1/d, & i = a, j \neq b \end{cases} \tag{40}$$

(where here, and below, all expectations are over the set of $d$ substitution effects on a single trait, $i$). Using eqs. 39-40 and eq. 2, we can now derive the mean and variance of the value of trait $i$.

$$\begin{aligned} E\left(z_i - o_i\right) &= z_{\mathrm{P1},i} - o_i + E\left(\sum_{j \in J_{hom}} m_{ij}\right) + \frac{1}{2}E\left(\sum_{j \in J_{het}} m_{ij}\right) \\ &= z_{\mathrm{P1},i} - o_i + dp_2 E\left(m_{ij}\right) + dp_{12}\frac{1}{2}E\left(m_{ij}\right) \\ &= z_{\mathrm{P1},i} - o_i + d\left(p_2 + \frac{p_{12}}{2}\right)\frac{z_{\mathrm{P2},i} - z_{\mathrm{P1},i}}{d} \\ &= (1-h)(z_{\mathrm{P1},i} - o_i) + h(z_{\mathrm{P2},i} - o_i) \end{aligned} \tag{41}$$

Similarly, for the variance, we find:

$$
\begin{aligned}
\mathrm{Var}(z_i - o_i) = \mathrm{Var}(z_i) =& \mathrm{Var}\left(\sum_{j\in J_{hom}} m_{ij}\right) \\
&+ \frac{1}{2}\mathrm{Var}\left(\sum_{j\in J_{het}} m_{ij}\right) \\
&+ 2\mathrm{Cov}\left(\sum_{j\in J_{hom}} m_{ij}, \frac{1}{2}\sum_{b\in J_{het}} m_{ib}\right) \\
\\
=& dp_2\mathrm{Var}(m_{ij}) + dp_2(dp_2-1)\mathrm{Cov}(m_{ij}, m_{ib}) \\
&+ dp_{12}\mathrm{Var}\left(\frac{1}{2}m_{ij}\right) + dp_{12}(dp_{12}-1)\mathrm{Cov}\left(\frac{1}{2}m_{ij}, \frac{1}{2}m_{ib}\right) \\
&+ 2dp_2dp_{12}\mathrm{Cov}\left(\frac{1}{2}m_{ij}, m_{ib}\right) \\
\\
=& dp_2(1-p_2) \\
&+ dp_{12}(1-p_{12})\frac{1}{4} \\
&- dp_2p_{12} \\
\\
=& d\left[h(1-h) - \frac{p_{12}}{4}\right] \qquad (42)
\end{aligned}
$$

The key results of eqs. 10-13, now follow from combining eqs. 41 and 42 with the definition of the distance metric (eq. 4), and the scaling factor (eq. 6). Equation 12 also uses the definition of cosine similarity:

$$
\rho = \frac{\langle \mathbf{z}_{\mathrm{P}1} - \mathbf{o}, \mathbf{z}_{\mathrm{P}2} - \mathbf{o}\rangle_\lambda}{\|\mathbf{z}_{\mathrm{P}1} - \mathbf{o}\|_\lambda \, \|\mathbf{z}_{\mathrm{P}2} - \mathbf{o}\|_\lambda} \equiv \frac{\sum \lambda_i (z_{\mathrm{P}1,i} - o_i)(z_{\mathrm{P}2,i} - o_i)}{\sqrt{\sum \lambda_i (z_{\mathrm{P}1,i} - o_i)^2}\sqrt{\sum \lambda_i (z_{\mathrm{P}2,i} - o_i)^2}} \qquad (43)
$$

In Figure S1, we also use a comparable result for the standard deviation of the scaled distance. This result depends much more strongly on the (assumed) normality of the trait values, $z_i$, and so a failure of its predictions is indicative of non-normal effects. Under normality, we have

$$
\mathrm{Var}\left(r^2\right) \approx \frac{16}{d^2\left(\sum_i \lambda_i\right)^2} 2\sum_i \lambda_i^2 \mathrm{Var}(z_i)\left(\mathrm{Var}(z_i) + 2E^2(z_i - o_i)\right) \qquad (44)
$$

which yields the following prediction for fully homozygous genotypes:

$$
\sqrt{\mathrm{Var}(r^2)\frac{\left(\sum_i \lambda_i\right)^2}{32\sum_j \lambda_j^2} - 2h(1-h)\frac{\sum_i \lambda_i E^2(z_i - o_i)}{d^2\sum_j \lambda_j^2}} = 4h(1-h), \qquad p_{12} = 0 \qquad (45)
$$

and this is the quantity plotted in the right-hand panels of Figure S1.

The same approach is used to derive the results with phenotypic dominance (eqs. 27-30). The result for $V_\delta$ (eq. 29) follows directly from the (strong) assumption that the dominance deviations - the $\delta_{ij}$ - show no covariance between traits, or with the homozygous effects - the $m_{ij}$. For the mean trait values, we have an extra term, describing the Brownian bridge between the global heterozygote and the midparent, and so eq. 41 is replaced by

$$E\left(z_i - o_i\right) = z_{\mathrm{P1},i} - o_i + p_2(z_{\mathrm{P2},i} - z_{\mathrm{P1},i}) + p_{12}\left(z_{\mathrm{H},i} - z_{\mathrm{P1},i}\right)$$

$$= z_{\mathrm{P1},i} - o_i + h(z_{\mathrm{P2},i} - z_{\mathrm{P1},i}) + p_{12}\left(z_{\mathrm{H},i} - \frac{z_{\mathrm{P1},i} + z_{\mathrm{P2},i}}{2}\right)$$

and so, with $z_{\mathrm{P},i} = (z_{\mathrm{P1},i} + z_{\mathrm{P2},i})/2$ denoting the midparental trait value, the squared mean is:

$$\begin{aligned} E^2\left(z_i - o_i\right) = &\left(z_{\mathrm{P1},i} - o_i\right)^2 + h^2\left(z_{\mathrm{P2},i} - z_{\mathrm{P1},i}\right)^2 + p_{12}^2\left(z_{\mathrm{H},i} - z_{\mathrm{P},i}\right)^2 \\ &+ 2h\left(z_{\mathrm{P1},i} - o_i\right)\left(z_{\mathrm{P2},i} - z_{\mathrm{P1},i}\right) \\ &+ 2p_{12}\left(z_{\mathrm{P1},i} - o_i\right)\left(z_{\mathrm{H},i} - z_{\mathrm{P},i}\right) \\ &+ 2hp_{12}\left(z_{\mathrm{P2},i} - z_{\mathrm{P1},i}\right)\left(z_{\mathrm{H},i} - z_{\mathrm{P},i}\right) \end{aligned}$$

After applying the scaling of eq. 6, and applying the cosine rule, this reduces to $M + M_\delta$ (eqs. 13 and 30).

#### Appendix 2

In this appendix, we briefly derive the maximum value of $r_{12}^2$. First, let us denote as $\mathbf{m}_i$ the $n$-dimensional vector of phenotypic change caused by the $i^{th}$ substitution differentiating the parental lines (and recalling that substitutions are defined relative the P1 parental genotype; see Figures 1 and 3). The magnitude of $\mathbf{m}_i$ is

$$\|\mathbf{m}_i\| = \|\mathbf{m}_i\|_\lambda \equiv \sqrt{\sum_{j=1}^{n} \lambda_j m_{ji}^2} \tag{46}$$

Note that here, and thoughout this appendix, we will drop the $\lambda$ subscript for brevity, but the $\lambda_j$ parameters are still used to capture differences between traits in the strength of selection, and the size of factors fixed (see Appendix 1).

Using this notation, the numerator of eq. 15, which is the phenotypic distance between the parents, is equal to the magnitude of the sum of the $\mathbf{m}_i$.

$$\|\mathbf{z}_{\mathrm{P2}} - \mathbf{z}_{\mathrm{P1}}\|^2 = \left\| \sum_{i=1}^{d} \mathbf{m}_i \right\|^2 \tag{47}$$

The denominator of eq. 15 is the "reference distance", defined as the expected length of a random walk in phenotype space, conditioned on the observed distribution of effect sizes. Under a random walk, the vectors will be orthogonal on average, and so, given the way that the $\lambda_i$ are defined (Appendix 1), this distance is equal to the sum of the squared magnitudes.

$$\left\| \sqrt{d}\mathbf{1} \right\|^2 = \sum_{i=1}^{d} \|\mathbf{m}_i\|^2$$

$$= d\left[\mathrm{Var}\left(\|\mathbf{m}_i\|\right) + E^2\left(\|\mathbf{m}_i\|\right)\right] \tag{48}$$

Finally, we note that the length of a sum of vectors is maximized when they all point in the same direction, so that the magnitude of the sum is the sum of the magnitudes. In this case, the maximised squared magnitude is

$$\max\left( \left\| \sum_{i=1}^{d} \mathbf{m}_i \right\|^2 \right) = \left( \sum_{i=1}^{d} \|\mathbf{m}_i\| \right)^2$$

$$= d^2 E^2\left(\|\mathbf{m}_i\|\right)$$

Putting these together, we have

$$\max\left(r_{12}^2\right) = 4\frac{\max\left(\left\|\sum_{i=1}^{d}\mathbf{m}_i\right\|^2\right)}{\sum_{i=1}^{d}\|\mathbf{m}_i\|^2}$$

$$= \frac{4d}{1+CV^2(\|\mathbf{m}_i\|)} \quad (49)$$

where $CV(\|\mathbf{m}_i\|) = \mathrm{Std}(\|\mathbf{m}_i\|)/E(\|\mathbf{m}_i\|)$ is the coefficient of variation in the magnitudes. When populations adapt to a distant stationary optimum via new mutations of variable sizes, (Orr, 1998b) showed that the $\|\mathbf{m}_i\|$ are often approximately exponentially distributed. In this case, $CV(\|\mathbf{m}_i\|) \approx 1$, so that $\max\left(r_{12}^2\right) \approx 2d$. The absolute maximum of $4d$ occurs when all vectors are of equal magnitude, such that $CV(\|\mathbf{m}_i\|) = 0$. This also follows from the triangle inequality.

### Appendix 3

In this Appendix, we give full details of the methods used for the simulations, whose results are presented in Figures 4-7, S1 and S3-S7.

##### A3.1 General simulation methods

We simulated the evolution of multi-locus diploid Wright-Fisher populations with selection. We considered two populations, each of $N$ individuals. Generations were discrete and non-overlapping, and individuals were treated as simultaneous hermaphrodites. Each generation, within each population, $2N$ parents were chosen with replacement, with probabilities proportional to their fitnesses. Gametes were generated from the parental genomes with recombination and mutation. Assumptions about recombination varied (see below). Each generation, the number of new mutations was drawn from a Poisson distribution, with mean $2NU$, and these were placed on the $2N$ gametes at random. The model assumed an effectively infinite number of sites, and so each new mutation took place at a unique, and randomly-chosen location in the genome, with no back mutation. Random numbers were generated using MTRand (Matsumoto and Nishimura, 1998). Simulations began with an ancestral population that was genetically uniform, although standing variation was allowed to accumulate during the divergence. For simulations with gene flow (scenarios 1-2), each of the $N$ offspring in a given population was formed from parents originating from the other population, with a probability $m$.

At the end of each simulation, we chose a single genotype from each population, which was then treated as one of the parental lines for hybridization. For allopatric runs, this genotype contained only the fixed mutations, while for parapatric runs we chose a genotype that contained all alleles whose frequencies differed by >50% across the two populations. In all cases, the phenotype associated with this genotype was found to be close to the modal phenotype of the polymorphic individuals in the population.

For all simulations, fitness was determined from eq. 5, with $\alpha = \frac{1}{2}$, and we assumed no differences between traits in the distribution of mutations, or in the strength of selection. We note that this does not preclude realised differences in the $\lambda_i$ parameters, because different traits might fix mutations with different distributions of sizes, especially if they experience different histories of environmental change (see below). To plot scaled distance values (eq. 6), we required an estimate of $\sum \lambda_i$, and this was taken from the sample variances of the realized $m_{ij}$. This would not work in the general case, but works acceptably well for our choice of fitness function, i.e., independent selection of unit strength on all traits ($A$ equal to the identity matrix, in the notation of eq. 33 from Appendix 1). All simulation code was written in C, and is available as online supplementary information.

##### A3.2 Description and justification of parameter values

As described in Table 1, simulations were run under a range of population genetic parameter values. In this section of the Appendix, we explain and justify our choices of these values.

The key population genetic parameters are the rates of recombination and mutation (Table 1). These determine the extent to which mutations will appear in a variety of genetic backgrounds during their fixation, and thus, the extent to which epistatic interactions might influence the divergence process. With this in mind, we compared very high and very low rates of recombination. In particular, we assumed either free recombination among all sites, or that all mutations were found on a single chromosome of map length 100cM with Haldane's mapping function. In other words, the number of crossover events was a Poisson random variable, with mean 1, and crossover locations were positioned at random. From Zeng (1990; see also eq. 9.3 of Lynch and Walsh, 1998), this corresponds to a mean crossover fraction of

$$\bar{c} = \frac{1}{2} - \frac{2L - 1 + e^{-2L}}{4L^2} = \frac{1 - e^{-2}}{4} \approx 0.216... \tag{50}$$

where $L = 1$ is the map length in Morgans. This value, which compares to $\bar{c} = \frac{1}{2}$ under free recombination, is lower than all of the empirical estimates compiled and reported by Lynch and Walsh, 1998 (Table 9.2).

For mutation rates, we chose values such that, when $N = 1000$, $2NU$ was as high as 10, generating high levels of segregating variation, and as low at 0.01, such that many mutations will reach fixation or loss, without appearing in a range of genetic backgrounds.

Other parameters affect the overall shape of the fitness landscape (Table 1). For the number of phenotypic traits, $n$, we chose to simulate under $n = 2$ and $n = 20$. As shown in a range of previous work, these values can lead to qualitatively different dynamics during divergence (e.g., Lourenço et al., 2011; Orr, 2000; Poon and Otto, 2000; Roze and Blanckaert, 2014; Zhang, 2012). In general, populations with $n = 20$ tend to have higher genetic loads, and a reduced ability to track moving environmental optima, while $n = 2$ maximizes the extent of stochastic correlations between trait values. The curvature of the fitness landscape, $k$, also has strong effects on the divergence (Fraïsse et al., 2016; Roze and Blanckaert, 2014). We chose the value $k = 2$, which leads to a Gaussian function. This is the most widely-studied version of the model, and has theoretical justification as a truncated Taylor expansion of an arbitrary function (Martin, 2014; Martin and Lenormand, 2006a), and some empirical support, from the distribution of new mutations (Martin et al., 2007; Martin and Lenormand, 2006b). The value $k = 2$ is also analytically tractable, since log fitness is directly proportional to the distance measure: $\ln w \propto \|\mathbf{z} - \mathbf{o}\|_\lambda^2$. We also chose $k = 6$, which yields a "table-like" fitness landscape, where small deviations from the optimum ($\|\mathbf{z} - \mathbf{o}\|_\lambda < 1$) all have similarly high fitness, while larger deviations ($\|\mathbf{z} - \mathbf{o}\|_\lambda > 1$), see a precipitous decline in fitness. This higher value of $k$ is better able to account for the very strong negative epistasis observed in some experimental studies of speciation genetics (Fraïsse et al., 2016).

Still other parameters specify the distribution of mutational effects on the phenotype. In all cases, we assumed that mutations showed universal pleiotropy (Martin, 2014; Sella and Barton, 2019). For the homozygous effects, we used two approaches: a "top-down" approach (Poon and Otto, 2000), where mutational vectors took a random direction in the $n$-dimensional space, and with magnitudes drawn from some known distribution. We also used a "bottom-up" approach, where the mutational effect on each trait was drawn from an i.i.d. normal distribution. The second approach tends to suppress mutations of small effect unless $n$ is very small (Orr, 2000; Wingreen et al., 2003), which acts to increase the possible upper bound on $r_{12}^2$ (see Appendix 2). Arguably, this also makes the approach biologically unrealistic with larger $n$ (Orr, 2000), although it remains widely used. For both top-down and bottom-up approaches, we chose parameters to fix the mean strength of selection against deleterious mutations appearing in an optimal genotype, which we denote $\bar{s}$. In particular, if we denote the mutational effect on trait $i$ as $x_i$, for the bottom-up approach, we assumed $x_i \sim N(0, \sigma^2)$, where

$$\sigma^2 = \frac{1}{2}\left(-\frac{\bar{s}}{\alpha}\frac{\Gamma(n/2)}{\Gamma((n+k)/2)}\right)^{2/k} \tag{51}$$

For the top-down approach, the magnitude of each mutation, $E\left(\sqrt{\sum_{i=1}^{n} x_i^2}\right) \equiv x$, was randomly generated, such that $x^{k/2}$ was exponentially distributed, and the mean of the exponential distribution was set at:

$$\lambda = \left(\frac{-\bar{s}}{\alpha k!}\right)^{1/k} \tag{52}$$

(Fraïsse et al., 2016). We also varied the mean strength of selection, $\bar{s}$, choosing values such that, when $N = 1000$, we had either $N\bar{s} = -10$, or $N\bar{s} = -0.1$. As such, selection was either effective on average, or ineffective on average.

Finally, for migration rates in the parapatric simulations (scenarios 1-2), we chose migration rates such that $Nm = 0.1$ and $Nm = 1$. Migration at these levels leads to different levels of differentiation under neutrality. We did not explore lower values, where dynamics tend to resemble allopatry, or larger values, which tend to prevent genomic divergence, due to allele swamping (Débarre et al., 2015; Dittmar et al., 2016; Griswold, 2006; Yeaman and Whitlock, 2011).

##### A3.3 Description of demographic/environmental scenarios

We simulated populations under different environmental/demographic scenarios, as summarized in Figure 4. To simulate an abruptly jumping optimum (labelled J in Figure 4), the trait values of the initial population were all placed at the origin (such that all $z_i = 0$), but the optimum value of one of the $n$ traits was set to 1. In consequence, the initial population had a fitness of $e^{-\alpha\|\mathbf{z}-\mathbf{o}\|_\lambda^k} = e^{-1/2} \approx 60\%$ relative to an optimal population. To simulate smooth, cyclical change (labelled C in Figure 4), for one of the $n$ traits, we set the optimum, at generation $t$ to:

$$\frac{1}{2}\left[1+\sin\left(\frac{2\pi t}{T}-\frac{\pi}{2}\right)\right]$$

This function oscillates over a period of $T$ generations between 0 and 1. We chose to set $T = 10^5$ for all parameter combinations. Because we simulated a model in which all $n$ traits were equivalent, apart from any changes in their optimum, the numbering and orientation of the traits was arbitrary. In the allopatric runs, this allowed us to relabel the traits in order to simulate moving optima on different traits, or to change the sign of all mutational effects, in order to simulate moving optima on the same trait in different directions. Population size remained stable in all simulations, except for scenario 13, where we included a transient bottleneck. To do this, we assumed that the population had a fixed size of $N = 100$ individuals for the first $25/U$ generations, while the next and all subsequent generations had $N = 1000$ offspring.

### Appendix 4

In this appendix, we briefly extend results for Fisher's model with two optima, to encompass hybrids scored in intermediate environments, or in patchy ecotones, where each individual experiences one of two alternative habitats, with some fixed probability. First, let us consider a patchy ecotone, in which hybrids experience habitat B with probability $q$, and habitat A with probability $1-q$. We begin by including the environmental indicator variable of Rundle and Whitlock (2001) in eq. 16, and write

$$\begin{aligned} E\left(r^2\right) =& r^2_{..} \\ &+\tfrac{1}{2}\theta_E\left(r^2_{.\mathrm{B}}-r^2_{.\mathrm{A}}\right) \\ &+\left(h-\tfrac{1}{2}\right)\left(r^2_{2.}-r^2_{1.}\right) \\ &+\tfrac{1}{2}\left(h-\tfrac{1}{2}\right)\theta_E\left(\left(r^2_{1\mathrm{A}}-r^2_{1\mathrm{B}}\right)-\left(r^2_{2\mathrm{A}}-r^2_{2\mathrm{B}}\right)\right) \\ &-p_{12} \\ &+4h(1-h)\left(1-\tfrac{1}{4}r^2_{12}\right) \end{aligned} \tag{53}$$

In this case, we can take expectations over the indicator variable in eq. 53, so that $E\left(\theta_E\right)=2q-1$. In this way, we find

$$\begin{aligned} E_{PE}\left(r^2\right) =& r^2_{.\mathrm{A}}+q\left(r^2_{.\mathrm{B}}-r^2_{.\mathrm{A}}\right) \\ &+\left(h-\tfrac{1}{2}\right)\left[r^2_{2\mathrm{A}}-r^2_{1\mathrm{A}}+q\left(r^2_{1\mathrm{A}}+r^2_{2\mathrm{B}}-r^2_{1\mathrm{B}}-r^2_{2\mathrm{A}}\right)\right] \\ &-p_{12}+4h(1-h)\left(1-\tfrac{1}{4}r^2_{12}\right) \end{aligned} \tag{54}$$

where the "$PE$" indicates a patchy ecotone. Equation 54 reduces to results for a single environment when $q=0$ or $q=1$. Now let us compare this result to predictions for a single environment, with an intermediate optimum. To model this, let us assume that the optimum is intermediate between two other optima, A and B, which might represent the parental habitats. Let us now treat $0 \leq q \leq 1$ as measure of the closeness of the intermediate optimum to optimum B. With this definition, we can write the optimum value for trait $i$ in the intermediate environment as

$$o_i = o_{\mathrm{A},i}+q(o_{\mathrm{B},i}-o_{\mathrm{A},i}) \tag{55}$$

where $o_{\mathrm{A},i}$ and $o_{\mathrm{B},i}$ are the optimum trait values in the two parental habitats. Using this definition in eq. 16, we find:

$$E_{IO}\left(r^2\right) = E_{PE}\left(r^2\right)-q(1-q)r^2_{\mathrm{AB}} \tag{56}$$

where "$IO$" denotes results with an intermediate optimum. Equations 54 and 56 differ only by the term $-q(1-q)r^2_{\mathrm{AB}}$. This implies that the additive effects, $\{\alpha_\mathrm{I}\}$, are identical for patchy ecotones, and intermediate habitats, if we use the appropriate definitions of $q$.
