## Supplementary Figures for "The geometry and genetics of hybridization"

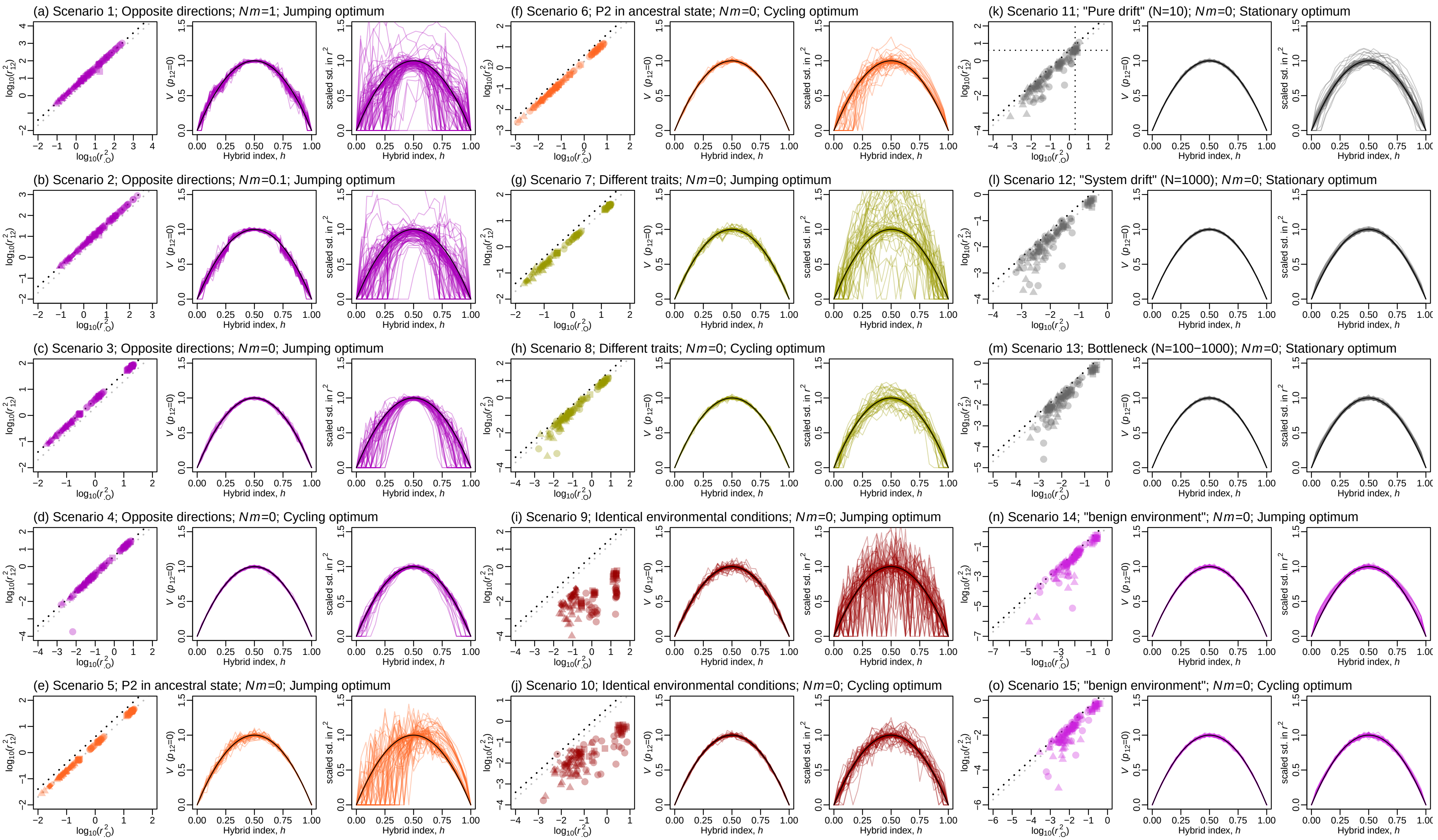
(a) Scenario 1; Opposite directions; Nm=1; Jumping optimum
(b) Scenario 2; Opposite directions; Nm=0.1; Jumping optimum
(c) Scenario 3; Opposite directions; Nm=0; Jumping optimum
(d) Scenario 4; Opposite directions; Nm=0; Cycling optimum
(e) Scenario 5; P2 in ancestral state; Nm=0; Jumping optimum
(f) Scenario 6; P2 in ancestral state; Nm=0; Cycling optimum
(g) Scenario 7; Different traits; Nm=0; Jumping optimum
(h) Scenario 8; Different traits; Nm=0; Cycling optimum
(i) Scenario 9; Identical environmental conditions; Nm=0; Jumping optimum
(j) Scenario 10; Identical environmental conditions; Nm=0; Cycling optimum
(k) Scenario 11; "Pure drift" (N=10); Nm=0; Stationary optimum
(l) Scenario 12; "System drift" (N=1000); Nm=0; Stationary optimum
(m) Scenario 13; Bottleneck (N=100−1000); Nm=0; Stationary optimum
(n) Scenario 14; "benign environment"; Nm=0; Jumping optimum
(o) Scenario 15; "benign environment"; Nm=0; Cycling optimum
$\log_{10}(r^2_{12})$
$\log_{10}(r^2_{\cdot O})$
$V\ (p_{12}=0)$
scaled sd. in $r^2$
Hybrid index, h

### Figure S1

Detailed presentation of the simulation results summarised in Figures 4-5. Each set of three panels shows results for 128 distinct combinations of population genetic parameters, under a single divergence scenario, as described in Table 1. Left-hand panels compare the quantities $r^2_{12}$ and $r^2_{.O} \equiv \frac{1}{2}\left(r^2_{1O} + r^2_{2O}\right)$ (eqs. 14-15). The black lines show the prediction $r^2_{12} = 4r^2_{.O}$ corresponding to $\varepsilon = 1$, and the grey line shows $r^2_{12} = 2r^2_{.O}$, corresponding to $\varepsilon = 1/2$ (eq. 19; Figure 2b). Plotting characters indicate the curvature of the fitness landscape, $k$ (eq. 5) and the number of phenotypic traits, $n$, that were used in the simulations: $k = 2$, $n = 2$ (circles), $k = 2$, $n = 20$ (squares); $k = 6$, $n = 2$ (triangles), $k = 6$, $n = 20$ (diamonds). However, there are few consistent differences between these regimes. The middle panels show $V$ (eq. 11) and match Figure 5a. Right-hand panels show corresponding predictions for the scaled standard deviation in $r^2$. These predictions depend strongly on the assumption of multivariate normality among the fixed effects, and are described in eq. 45 of Appendix 1. As expected (Orr, 1998b), panels show that deviations from normality are most severe when divergence takes place via natural selection (scenarios 1-10), and especially with abruptly changing optima (scenarios 1-3, 5, 7 and 9). Nevertheless, despite these highly non-normal distributions, predictions for $V$ (middle panels) remain reasonably accurate. Also of note is the results for divergence via "pure drift" (scenario 11). In this case, several parameter combinations yielded results that were close to the expectations of $E\left(r^2_{.O}\right) = 2$ and $E\left(r^2_{12}\right) = 4$, if the parental phenotypes had both wandered around half of the "reference distance" from the optimum, via unconstrained random walks (Simon et al., 2018; see also Figure S5). Other parameter combinations yielded much smaller values, indicating that selection became effective once the population had drifted sufficiently far from its optimum.

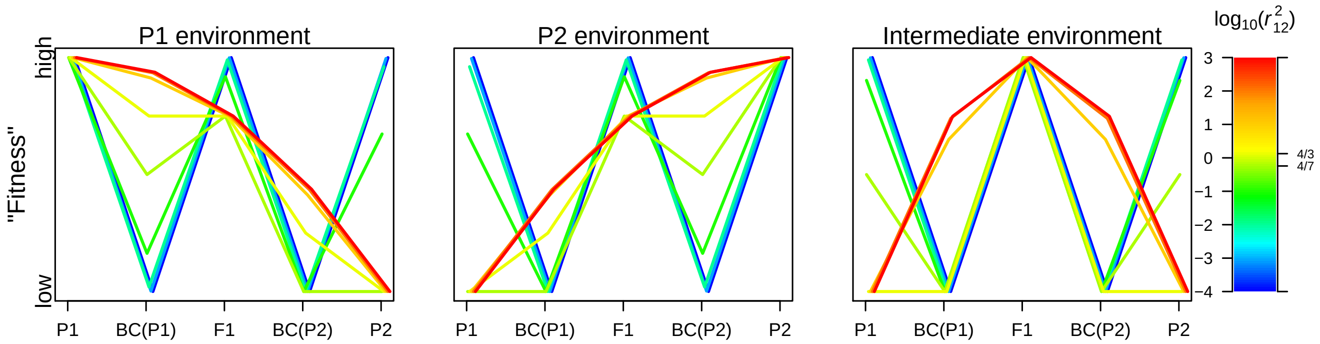


#### Figure S2

An illustration of the predictions of Fisher's geometric model in a situation of local adaptation, where two populations are each well adapted to a distinct environmental optimum. In this case, results depend solely on the scaled distance between the parental phenotypes, $r_{12}^2$. When $r_{12}^2$ is large, results show a characteristic pattern of ecological isolation in the parental habitats, and hybrid advantage in intermediate habitats. When $r_{12}^2$ becomes small, results in all three habitats approach the same "W-shaped" pattern of intrinsic isolation. These two extremes are illustrated in Figure 1 of Rundle and Whitlock (2001). The figure also shows the inflection points at $r_{12}^2 = 4/3$ (which leads to equal fitnesses for the fitter backcross and the F1), and at $r_{12}^2 = 4/7$ (which leads to equal fitnesses for the less fit parent and the less fit backcross). All results shown use eq. 16, with $M = h^2 r_{12}^2$, $M = (1-h)^2 r_{12}^2$, and $M = [\frac{1}{4} - h(1-h)h]r_{12}^2$ for the P1, P2 and intermediate environment respectively, and assuming $\mathrm{Var}(h) = 0$. Results are plotted on an arbitrary scale, to represent the rank order of the fitness values. The details of the crosses are given in legend to Figure 6, where the predictions are compared to simulations.

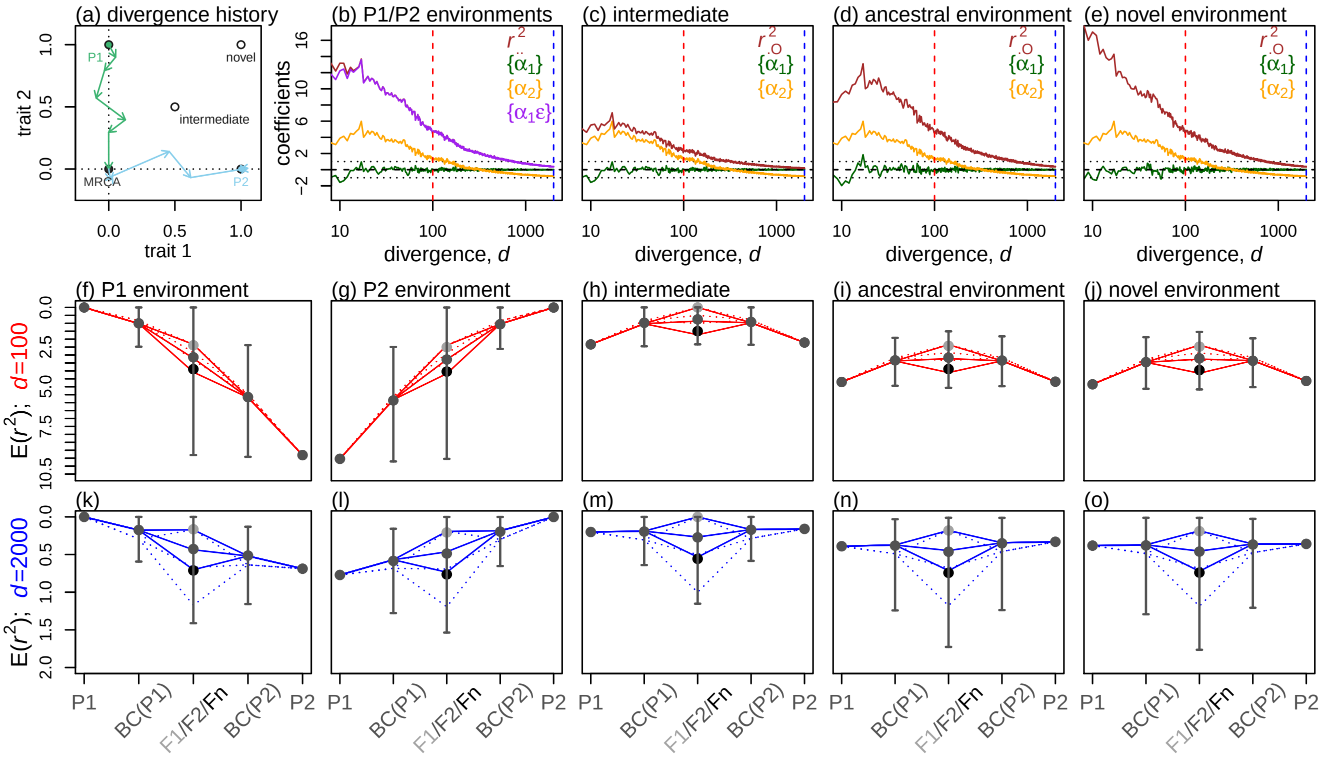


### Figure S3

Hybridization with local adaptation under Fisher's geometrical model, with low recombination. All details match Figure 6, except for the levels of recombination assumed during the simulations. While Figure 6 assumed free recombination among all loci, here, during both divergence and hybridization, we assumed a single chromosome of map length 100cM and Haldane's mapping function. This corresponds to an average recombination rate of $\bar{c} \approx 0.216$ between randomly chose pairs of sites (see Appendix 3). Analytical predictions used eq. 16 with $\mathrm{Var}(h) = (1-2\bar{c})/4$ for the homozygous "Fn" (produced by amphimictic selfing among F1 gametes), $\mathrm{Var}(h) = (1-2\bar{c})/8$ for the F2, and $\mathrm{Var}(h) = (1-2\bar{c})/16$ for the backcrosses. For comparison, analytical predictions with free recombination are shown as dotted lines.

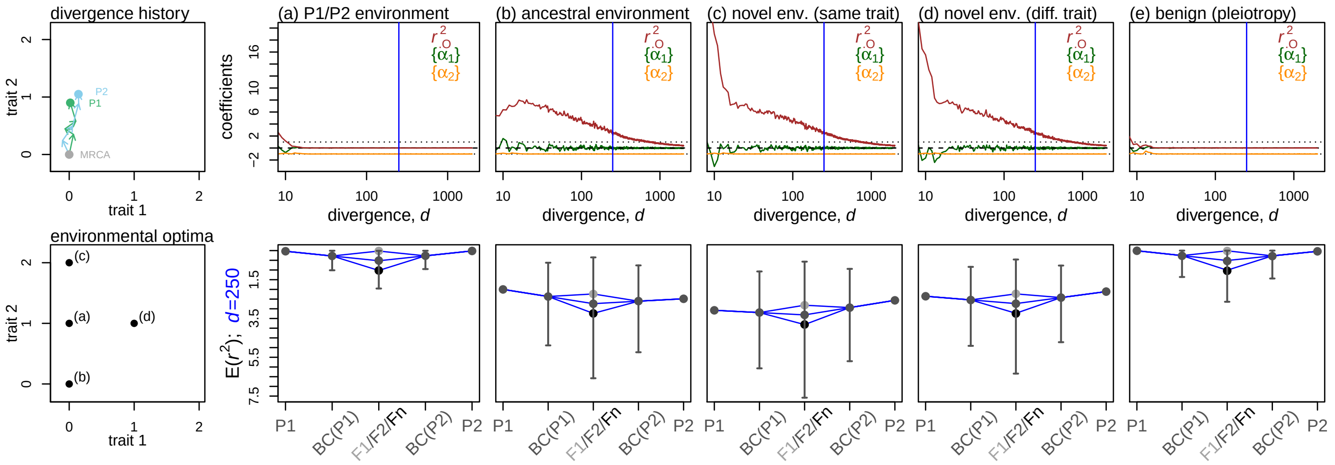


### Figure S4

Hybridization, under Fisher's geometrical model, when parental populations have adapted, independently, to the same identically shifting optima (this is shown, in cartoon, in the upper left-hand panel, and corresponds to "scenario 9" of Figure 4). Hybrids were scored in four distinct environments, as indicated in the lower left-hand panel, and with results shown in panels (a)-(d). Panel (e) shows results in a "benign" environment, where the trait with the shifting optimum no longer affected fitness (as in scenarios 14-15 in Figure 4). In these cases, results depended solely on pleiotropic effects of substitutions affecting the remaining trait, where the optimum remained stationary. In all cases, results show convergence to a characteristic pattern of intrinsic isolation (eq. 20). All other details match Figure 6.

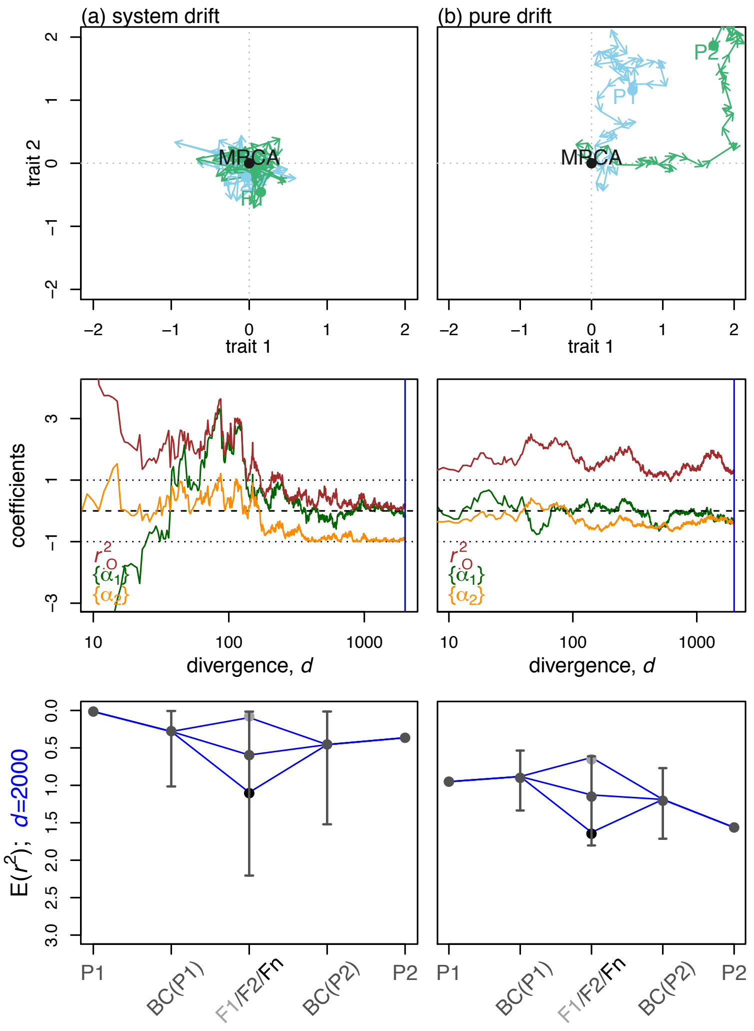


### Figure S5

Hybridization, under Fisher's geometrical model, when parental populations diverged via drift (i.e., variants of scenarios 11-12 in Figure 4). (a) With "system drift", the populations remained close to the optimum, but evolved along nearly-neutral pathways in the fitness landscape. This was achieved by repeating simulations with $N = 100$, $N\bar{s} = -0.01$ and $n = 2$. Results show that phenotypes move stochastically, but returning to values predicted by the characteristic pattern of intrinsic isolation (eq. 20). (b) With "pure drift", selection was so ineffective that populations continued to accrue mutations regardless of their effects. This was achieved by reducing the population size ($N = 50$ such that $N\bar{s} = -0.005$), and increasing the number of traits ($n = 20$). In this case, the evolution of the traits resembles a random walk, with $E\left(r^2_{\cdot O}\right) = 2$. Because the divergence is largely deleterious in both lines, the fitness of hybrids increases with their heterozygosity, in agreement with predictions from the classical theory of inbreeding (Simon et al., 2018; Wright, 1922). All other details match Figure 6.

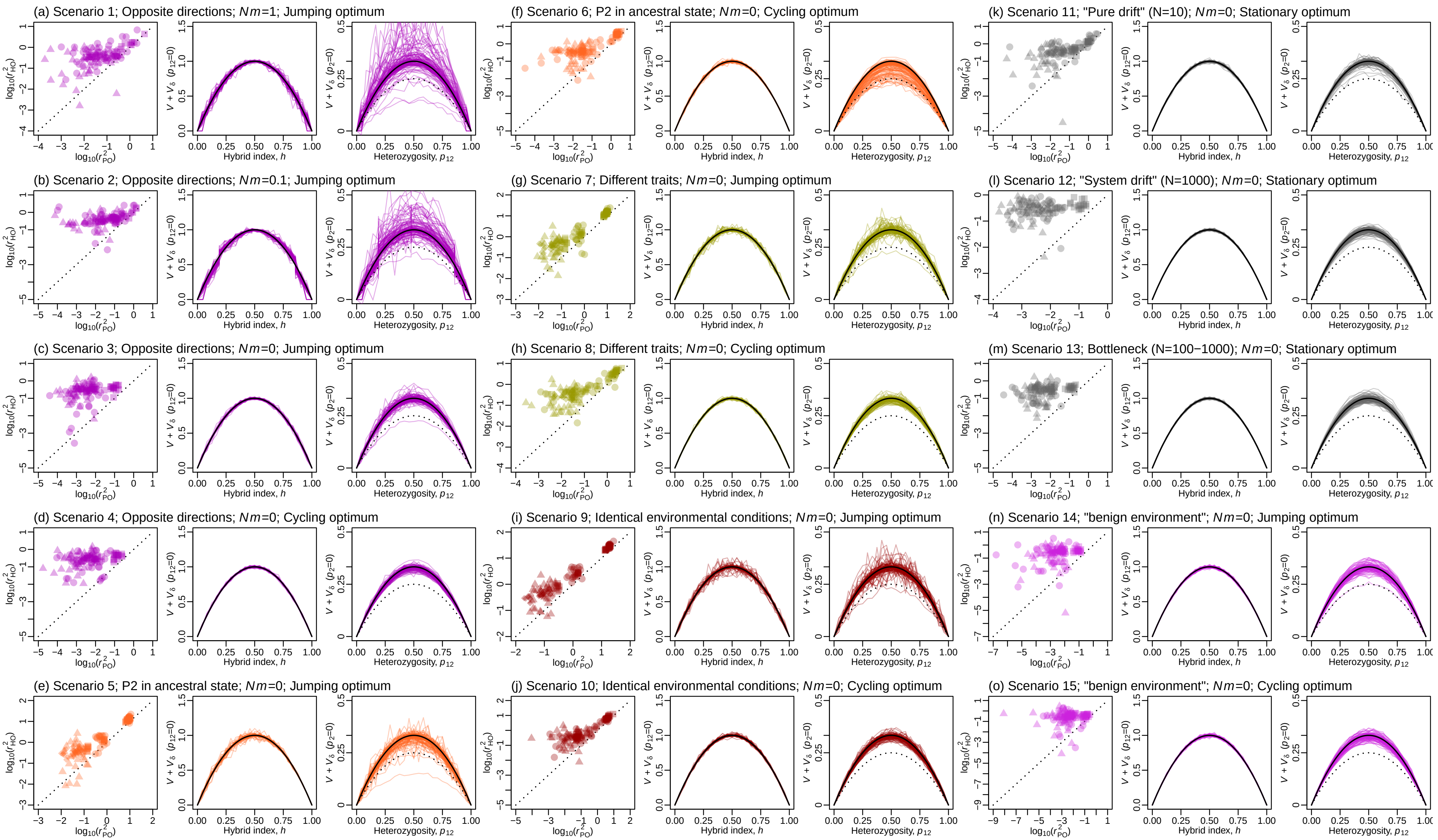
(a) Scenario 1; Opposite directions; Nm=1; Jumping optimum
(b) Scenario 2; Opposite directions; Nm=0.1; Jumping optimum
(c) Scenario 3; Opposite directions; Nm=0; Jumping optimum
(d) Scenario 4; Opposite directions; Nm=0; Cycling optimum
(e) Scenario 5; P2 in ancestral state; Nm=0; Jumping optimum
(f) Scenario 6; P2 in ancestral state; Nm=0; Cycling optimum
(g) Scenario 7; Different traits; Nm=0; Jumping optimum
(h) Scenario 8; Different traits; Nm=0; Cycling optimum
(i) Scenario 9; Identical environmental conditions; Nm=0; Jumping optimum
(j) Scenario 10; Identical environmental conditions; Nm=0; Cycling optimum
(k) Scenario 11; "Pure drift" (N=10); Nm=0; Stationary optimum
(l) Scenario 12; "System drift" (N=1000); Nm=0; Stationary optimum
(m) Scenario 13; Bottleneck (N=100−1000); Nm=0; Stationary optimum
(n) Scenario 14; "benign environment"; Nm=0; Jumping optimum
(o) Scenario 15; "benign environment"; Nm=0; Cycling optimum
Hybrid index, h
Heterozygosity, p12

#### Figure S6

Detailed presentation of the simulation results summarised in Figure 5c-f, which incorporate variable phenotypic dominance. Each set of three panels shows results for 128 distinct combinations of population genetic parameters, as described in Table 1, under a single divergence scenario (Figure 4). The left-hand panels compare the scaled distances to the optimum of the midparental phenotype (x-axis) and the global heterozygote (y-axis). There is a clear tendency for the global heterozygote to show lower fitness, allowing for a reduction in the fitness of F1 for genetically divergent parents. The middle panels of each set show the curves shown in Figure 5c, and confirm that the basic Brownian bridge approximation continues to apply well to homozygous hybrids. The right-hand panels in each set correspond to Figure 5e, and show the Brownian bridges formed by adding P2 alleles in heterozygous state to a P1 background. The black dotted lines show the additive prediction of $p_{12}(1-p_{12})$, and the black solid lines show $(1+4v)p_{12}(1-p_{12})$ with $v=1/12$ (corresponding the variance of a uniform distribution). A poor fit to the predictions is shown for the parapatric scenarios 1-2, when the divergence $d$ was very low ($d<50$). Also notable is the handful of runs for which the variance associated with heterozygous effects was lower than the expectations under additivity (naively implying $v<0$). These runs all involved abruptly shifting optima (i.e., scenarios 1-3, 7 and 9), and/or scenarios where all of the evolution took place in P1 (scenarios 5-6). In these runs, one (but not both) of the populations fixed several large-effect dominant mutations, and this led to strong directional dominance, and low variance in the heterozygous effects of the P2 alleles. As such, they imply covariance between homozygous and dominance effects, that was not included in our analytical treatment. All other plotting details match Figure S1.

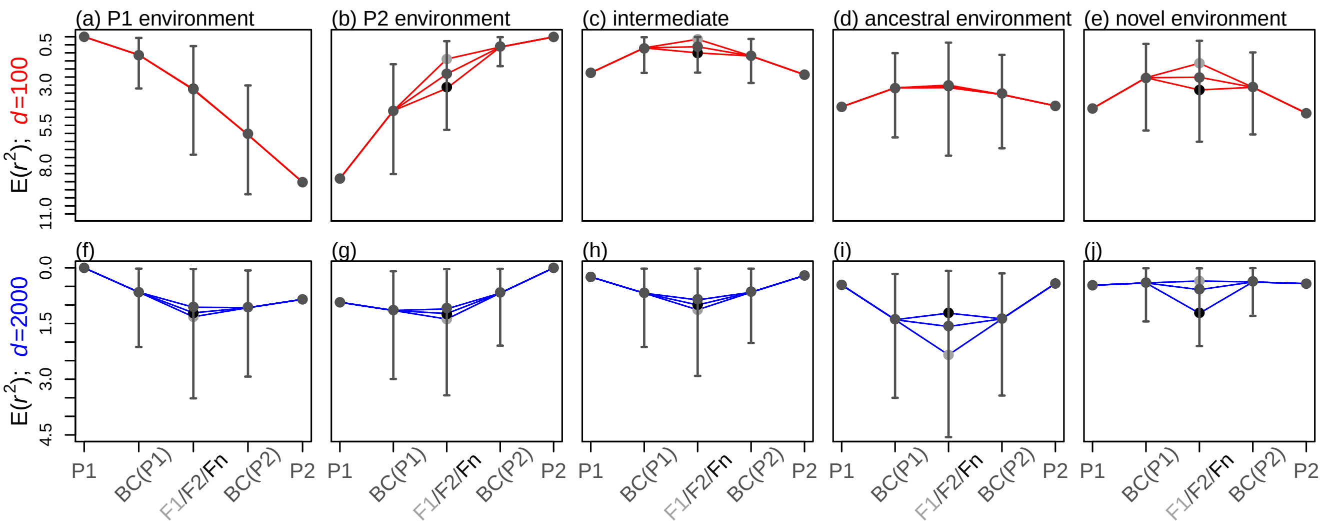


#### Figure S7

Hybridization under Fisher's geometrical model with local adaptation, and variable phenotypic dominance. All details match the lower panels of Figure 6, except that, during the process of divergence, simulated mutations had variable dominance effects. In particular, the heterozygous effect of each mutation was generated by multiplying its homozygous effect by a uniformly distributed random number. The qualitative changes introduced by phenotypic dominance are clearly visible. First, the F1 has reduced fitness at high levels of divergence (compare, for example, S7i to Figure 6n). Second, the effects of heterozygosity are now environment dependent (extrinsic); this is clear from comparing the middle crosses (F1/F2/Fn) in panels a and b, or d and e. Third, because of "directional dominance", the reciprocal backcrosses may have different fitnesses, even when the parents have identical fitnesses (these effect is evident, albeit weakly, in panels c-e). The red and blue lines used eqs. 28-30 with the simulated phenotypes of the parents and global heterozygote, and setting $v = 1/12$.
